# The dorsal visual pathway represents object-centered spatial relations for object recognition

**DOI:** 10.1101/2021.11.12.468414

**Authors:** Vladislav Ayzenberg, Marlene Behrmann

## Abstract

Although there is mounting evidence that input from the dorsal visual pathway is crucial for object processes in the ventral pathway, the specific functional contributions of dorsal cortex to these processes remain poorly understood. Here, we hypothesized that dorsal cortex computes the spatial relations among an object’s parts – a processes crucial for forming global shape percepts – and transmits this information to the ventral pathway to support object categorization. Using fMRI with human participants (females and males), we discovered regions in the intraparietal sulcus (IPS) that were selectively involved in computing object-centered part relations. These regions exhibited task-dependent functional and effective connectivity with ventral cortex, and were distinct from other dorsal regions, such as those representing allocentric relations, 3D shape, and tools. In a subsequent experiment, we found that the multivariate response of posterior IPS, defined on the basis of part-relations, could be used to decode object category at levels comparable to ventral object regions. Moreover, mediation and multivariate effective connectivity analyses further suggested that IPS may account for representations of part relations in the ventral pathway. Together, our results highlight specific contributions of the dorsal visual pathway to object recognition. We suggest that dorsal cortex is a crucial source of input to the ventral pathway and may support the ability to categorize objects on the basis of global shape.

**Significance Statement:** Humans categorize novel objects rapidly and effortlessly. Such categorization is achieved by representing an object’s global shape structure, that is, the relations among object parts. Yet, despite their importance, it is unclear how part relations are represented neurally. Here, we hypothesized that object-centered part relations may be computed by the dorsal visual pathway, which is typically implicated in visuospatial processing. Using fMRI, we identified regions selective for the part relations in dorsal cortex. We found that these regions can support object categorization, and even mediate representations of part relations in the ventral pathway, the region typically thought to support object categorization. Together, these findings shed light on the broader network of brain regions that support object categorization.

## Introduction

A central organizing principle of the brain is that the visual system is segregated into a ventral visual pathway for recognizing objects and a dorsal visual pathway for locating and interacting with objects (Mishkin et al., 1983; Ungerleider & Haxby, 1994). However, research increasingly shows that the dorsal pathway computes some of the same object properties as the ventral pathway (Farivar, 2009; Freud et al., 2020; Freud et al., 2016), and may even play a functional role in object recognition (Freud et al., 2020; Holler et al., 2019). Despite these findings, the dorsal pathway is rarely included in conceptual or computational models of visual recognition (Gauthier & Tarr, 2016; Zhuang et al., 2021). Indeed, artificial neural network models (ANNs) trained for object recognition are almost exclusively modelled on ventral cortex processes (Blauch et al., 2021; Kubilius et al., 2019). One potential reason for this exclusion, is that the specific functional contributions of the dorsal pathway to object recognition are poorly understood.

The primary function of the dorsal pathway has long been considered to be the computation of visuospatial information in the service of coordinating actions (Goodale & Milner, 1992; Mishkin et al., 1983). However, dorsal cortex, particularly the posterior parietal cortex (PPC), also computes object properties relevant for recognition. For instance, many studies find robust sensitivity to shape information in the PPC (Bracci & Op de Beeck, 2016; Freud et al., 2017; Georgieva et al., 2008), akin to ventral object regions such as the lateral occipital complex (LOC; Grill-Spector et al., 2001; Kourtzi & Kanwisher, 2001). As in LOC, dorsal shape representations are seemingly robust to changes in size and orientation, as well as format (i.e., 3D vs. 2D; Konen & Kastner, 2008; Vaziri-Pashkam & Xu, 2019). Object representations in the dorsal pathway also appear to be relatively abstract, such that the multivariate responses in PPC corresponds to perceived semantic similarity among objects, even when controlling for low-level visual properties (Bracci & Op de Beeck, 2016; Jeong & Xu, 2016).

Although these studies highlight the similarities between dorsal and ventral pathways, object representations in dorsal cortex are not simply redundant with those in the ventral cortex (Bracci & Op de Beeck, 2016; Freud et al., 2015; Vaziri-Pashkam & Xu, 2019). What, then, are the unique contributions of the dorsal pathway to object recognition? One possibility, consistent with its role in visuospatial processing (Kravitz et al., 2011; Mishkin et al., 1983), is that dorsal cortex computes the *spatial relations* among an object’s component parts – that is, the object’s topological structure, but not the form of object parts themselves – and then propagates this information to the ventral pathway to support object recognition.

Many studies have demonstrated that a description of part relations is crucial for forming invariant ‘global shape’ representations (Biederman, 1987; Hummel, 2000), which may be key for recognizing objects across variations in viewpoint or across category exemplars (Ayzenberg & Lourenco, 2019; Hummel & Stankiewicz, 1996). Indeed, an inability to represent the part relations results in marked deficits in object recognition (Behrmann et al., 2006). Such a representation may be particularly important for basic-level object categorization because members of a category typically have similar spatial structures, but vary in regards to their component parts (Ayzenberg & Lourenco, 2019; Barenholtz & Tarr, 2006; Rosch et al., 1976).

Surprisingly, few studies have investigated whether the dorsal pathway represents object-centered part relations, with most, historically, focusing on allocentric spatial coding (Haxby et al., 1991), and even fewer have examined the relation between such coding in the dorsal pathway and object recognition processes in the ventral pathway (c.f. Zachariou et al., 2017). Thus, in the current study, we tested whether the dorsal visual pathway represents the relations among component parts and whether this information may support object recognition processes in the ventral pathway.

To this end, in a first experiment, we tested whether regions of dorsal cortex exhibit selectivity for part relations, and examined the extent to which coding in these regions are independent of allocentric relations and other object properties represented by the dorsal pathway, such as 3D shape and tools. We also examined whether regions that represent part relations exhibit task-dependent functional connectivity with ventral cortex. We used effective connectivity analyses to test the directionality of these interactions, and, specifically, whether dorsal cortex predicts the response of ventral cortex, rather than the other way around. In a second experiment, we investigate whether these dorsal regions can support object categorization and whether they do so by representing the relations among parts. Using a decoding approach we measured the ability of dorsal regions to classify naturalistic objects, and tested whether their response profile to these objects was best characterized by a computational model that computes that spatial relations among parts. Finally, as in Experiment 1, we examined the degree to which dorsal and ventral cortex interact during object perception, as well as the directionality of their interactions.

## Materials and Methods

### Participants

Sample sizes and procedures for Experiment 1 (https://aspredicted.org/WSV_W7L) and Experiment 2 (https://aspredicted.org/49C_D4C) were preregistered following pilot testing. We recruited 12 participants (3 female, 9 male; *M*_*age*_ = 27.50, SD = 3.61) for Experiment 1, in which functional regions of interest (ROIs) were identified, and 12 participants (6 female, 6 male; *M*_*age*_ = 26.83, SD = 3.7) for Experiment 2, in which the ROIs’ contributions to object recognition were explored. Where possible, the same participants completed both Experiment 1 and 2, so that their pre-defined functional ROIs could be used for analysis. In total, eight participants from Experiment 1 also participated in Experiment 2. The four new participants in Experiment 2 were scanned in a second session (following the scanning procedure of Experiment 1).

Sample sizes were determined on the basis of prior studies which typically recruited between 10 and 15 participants (e.g., Bracci & Op de Beeck, 2016; Freud et al., 2017; Jeong & Xu, 2016). Nevertheless, to ensure that our chosen sample size did not influence the results, all analyses were replicated with a larger sample. Specifically, for Experiment 1, we included data from the four new participants (3 female; 1 male) initially tested for Experiment 2 and scanned two more participants (1 female; 1 male) thereby bringing the total sample size to 18 participants. For Experiment 2, we scanned two additional participants (1 female; 1 male), bringing the total sample size to 14. However, in keeping with the spirit of open science practices, we focus our analyses on the preregistered sample sizes.

All participants were right-handed and had normal or corrected-to-normal visual acuity. Participants were recruited from the Carnegie Mellon University community, gave informed consent according to a protocol approved by the Institutional Review Board (IRB), and received payment for their participation.

## Experimental Design and Statistical Analysis

### MRI scan parameters and analysis

Scanning was done on a 3T Siemens Prisma scanner at the CMU-Pitt Brain Imaging Data Generation & Education (BRIDGE) Center. Whole-brain functional images were acquired using a 64-channel head matrix coil and a gradient echo single-shot echoplanar imaging sequence. The acquisition protocol for each functional run consisted of 48 slices, repetition time = 1 s; echo time = 30 ms; flip angle = 64°; voxel size = 3 × 3 × 3 mm. Whole-brain, high-resolution T1-weighted anatomical images (repetition time = 2300 ms; echo time = 2.03 ms; voxel size = 1 × 1 ×1 mm) were also acquired for each participant for registration of the functional images.

All images were skull-stripped (Smith, 2002) and registered to the Montreal Neurological Institute (MNI) 2mm standard template. Prior to statistical analyses, images were motion corrected, de-trended, and intensity normalized. To facilitate functional and effective connectivity analyses, 18 additional motion regressors generated by FSL were also included. All data were fit with a general linear model consisting of covariates that were convolved with a double-gamma function to approximate the hemodynamic response function. Data used to define regions of interest (ROIs) was spatially smoothed using a 6 mm Gaussian kernel. All other data were unsmoothed. All data were analyzed using the peak 100 voxels within a region (as defined by the functional localizer) or using a 6mm sphere (∼120 voxels) centered on the peak voxel. Qualitatively similar results were found for all analyses when ROI sizes were varied parametrically from 100 to 400 voxels (the size of the smallest ROI). Analyses were conducted using FSL (Smith et al., 2004), and the nilearn, nibabel, and Brainiak packages for in Python (Abraham et al., 2014; Kumar et al., 2020).

### Experiment 1: Localization of object-centered part relations

Participants completed four localizer scans to measure voxels activated by object-centered part relations, allocentric relations, 3D shape, and tools. The allocentric relations localizer was included to test whether ROIs are sensitive to part relations specifically, or to spatial relations more generally. Although dorsal regions are sensitive to many spatial properties (e.g., orientation), we chose to measure allocentric relations because of their conceptual similarity to object-centered part relations. Similarly, the 3D shape localizer was included to test whether these ROIs are sensitive to shape information as defined by part relations, or by shape properties more generally. We specifically chose to test 3D shape because extensive research has shown that dorsal cortex is particularly sensitive to the depth properties of objects (Gillebert et al., 2015; Van Dromme et al., 2016), and may transmit this information to ventral cortex to support recognition (Freud et al., 2020). Finally, the tool localizer was included to test whether ROIs that represent part relations do so for objects more generally, or exclusively for objects that afford action.

We used a ROI approach to define regions in parietal cortex that represent part relations. Then, we used independent data to test the selectivity of these ROIs to part relations or to other visual properties represented by the dorsal pathway, namely allocentric relations (Haxby et al., 1991), 3D shape (Georgieva et al., 2008), and tools (Mahon et al., 2007). Furthermore, we conducted conjunction analyses to examine the degree of overlap between dorsal ROIs sensitive to part relations and the other dorsal properties (allocentric relations, 3D shape, tools). Finally, we conducted task-dependent functional and effective connectivity analyses to examine the degree to which dorsal ROIs sensitive to part relations are correlated with ventral regions, and whether part-relation coding in dorsal ROIs precedes, and even predicts, object processing in ventral ROIs.

For each localizer, we defined posterior and anterior parietal ROIs by overlaying posterior intraparietal sulcus (pIPS) and anterior IPS (aIPS) binary masks and selecting voxels within those masks that survived a whole-brain cluster-corrected threshold (*p* < .001). Broad pIPS and aIPS masks were created by combining IPS0 with IPS1 and IPS2with IPS3 probabilistic masks, respectively, from the Wang et al. (2014) atlas. For comparison of the activation profiles from dorsal regions, an object-selective ROI in the ventral stream was defined similarly within the lateral occipital complex (LOC) probabilistic parcel (Julian et al., 2012).

### Object-centered part relations localizer

Participants completed six runs (320 s each) of an object-centered part relations localizer consisting of blocks of object images in which either *the spatial arrangement* of component parts varied from image to image (part-relations condition), while the parts themselves stayed the same; or the *features* of the component parts varied from image to image (feature condition), while the spatial arrangement of the parts stayed the same (Figure 1A). Objects could have one of 10 possible spatial arrangements, and one of 10 possible part features. Spatial arrangements were selected to be qualitatively different from one another as outlined by the recognition-by-components (RBC) model (e.g., end-to-end; end-to-middle; Biederman, 1987). The component parts were comprised of qualitatively different features as outlined by the RBC model (e.g., sphere, cube). Because many dorsal regions are particularly sensitive to an object’s orientation and axis of elongation (Sakata et al., 1998), objects were presented in the same orientations and were organized around the same elongated segment, ensuring they have identical principal axes. Stimuli subtended ∼6° visual angle on screen.

**Figure 1.**
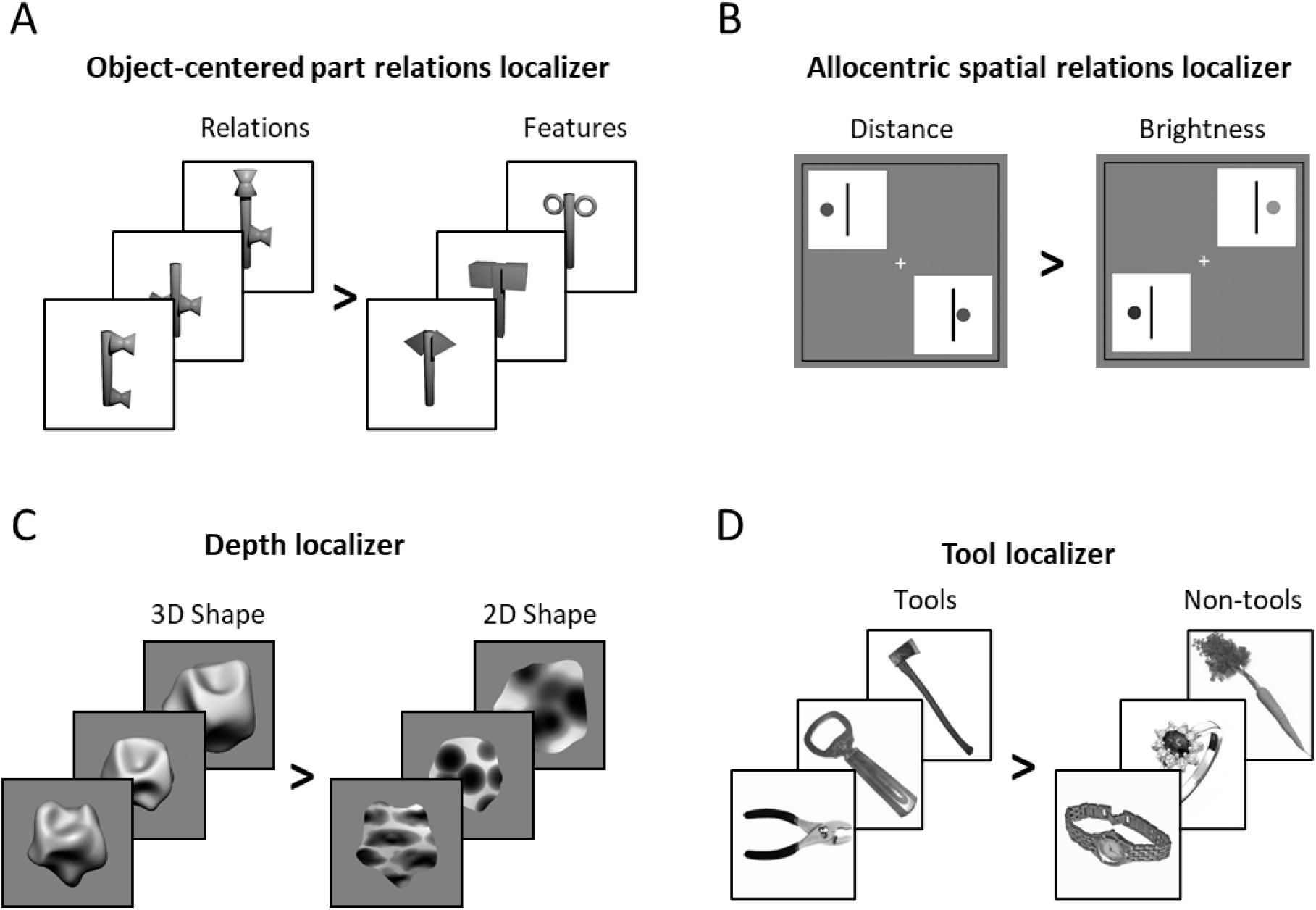
Example stimuli from the (A) object-centered part relations, (B) allocentric relations (C) depth, (D) and tool localizers used in Experiment 1.

Each block of the part relations localizer contained 20 images, displaying each spatial arrangement or part feature twice per block depending on the condition. Each image was presented for 800 ms with a 200 ms interstimulus interval (ISI) for a total of 20 s per block. To minimize visual adaptation, the location of object images on the screen varied by ∼2° every trial. The image order within the block was randomized. Participants also viewed blocks of a fixation cross (20 s). Participants viewed 5 repetitions of each block per run, with blocks presented in a pseudorandom order under the constraint that all three block types (relations, feature, fixation) were presented once before repetition. To maintain attention, participants performed an orthogonal one-back task, in which they responded via key press when detecting the repetition of an image on consecutive presentations.

Object-centered part relations ROIs in pIPS and aIPS were defined in each individual using 4 out of the 6 MRI runs as those voxels that responded more to the part-relations than the feature condition. Selectivity was measured for each voxel in an ROI by extracting standardized parameter estimates for each condition (relative to fixation) in left out runs (2 out of 6).

### Allocentric relations localizer

Participants completed two runs (368 s each) of the allocentric relations localizer wherein some blocks they judged whether displayed objects had the same allocentric relations, in this case the same distances between objects (distance condition), or had the same brightness (brightness condition; Zachariou et al., 2017). A nearly identical display was shown in both conditions, consisting of two diagonally arranged displays, each containing a line and circle (Figure 1B). In the distance condition, the allocentric relations (i.e., distances) between the line and circle, either matched across the two displays or differed. In the brightness condition, the brightness of the circles across the two displays either matched or differed. On each trial, participants were required to indicate whether the two displays were the same or different (according to distance or brightness). Each display subtended ∼4° visual angle on screen. Prior to the start of the scan, participants’ individual sensitivity to distance and brightness (blocked) was measured using an adaptive task where the distances and brightness of the stimuli was titrated until accuracy on each of the tasks was approximately 75%. We specifically used this allocentric localizer task because it has been well validated in human neuroimaging studies (Haxby et al., 1991; Zachariou et al., 2014).

Each block contained 10 distance or brightness trials, in which five trials had matching displays and five trials had different displays. Each trial was presented for 1700 ms with a 300 ms interstimulus interval (ISI) for a total of 20 s per block. The trial order within the block was randomized. Participants also viewed blocks of fixation (20s). Participants viewed 6 repetitions of each block per run, with blocks presented in a pseudorandom order under the constraint that all three block types (distance, brightness, fixation) were presented once before repetition.

Allocentric relation ROIs were defined in each individual as those voxels that responded more to the distance than the brightness condition. Selectivity was measured for each voxel in an ROI by extracting standardized parameter estimates for each condition (relative to fixation).

### Depth localizer

Participants completed two runs (308 s each) of a depth localizer wherein they viewed blocks of object images that contained 3D shapes as defined from depth shading cues (3D condition), or 2D shapes with comparable low-level properties (2D condition; Figure 1C). Each condition was comprised of ten 3D or 2D object images from Georgieva et al. (2008). All stimuli were ∼6° visual angle on screen. Each block contained 20 images, displaying each possible 3D or 2D image twice per block. Each image was presented for 700 ms with a 100 ms interstimulus interval (ISI) for a total of 16 s per block. The image order within the block was randomized. Participants also viewed blocks of fixation (16 s). Participants viewed 6 repetitions of each block per run, with blocks presented in a pseudorandom order under the constraint that all three block types (3D, 2D, fixation) were presented once before repetition. To maintain attention, participants performed an orthogonal one-back task, responding to the repetition of an image on consecutive presentations.

Depth ROIs were defined in each individual as those voxels that responded more to the 3D than the 2D condition. Selectivity was measured for each voxel in an ROI by extracting standardized parameter estimates for each condition (relative to fixation) in left out runs.

### Tool and object localizer

Participants completed two runs (340 s) of a tool localizer wherein they viewed blocks of object images that contained tools (tool condition), manipulable non-tool objects (non-tool condition), or box-scrambled object images (scrambled conditions; Figure 1D). Following previous work (Mahon et al., 2007), we define tools here as manipulable objects whose physical form is directly related to their function (e.g., a hammer). By contrast, manipulable non-tool objects are those that can be arbitrarily manipulated, but whose form is not directly related to their function (e.g., a carrot). Each condition was comprised of ten instances each of tools, non-tools, or scrambled object images from (Chen et al., 2018; Chen et al., 2016). Each block contained 20 images, displaying each possible tool, non-tool, or scrambled image twice per block. All stimuli subtended ∼6° visual angle on screen. Each image was presented for 700 ms with a 100 ms interstimulus interval (ISI) for a total of 16 s per block. The image order within the block was randomized. Participants also viewed blocks of fixation (16 s). Participants viewed 5 repetitions of each block per run, with blocks presented in a pseudorandom order under the constraint that all four block types (tool, non-tool, scrambled, fixation) were presented once before repetition. To maintain attention, participants performed an orthogonal one-back task, responding to the repetition of an image on consecutive presentations.

Tool ROIs were defined in each individual as those voxels that responded more to the tool than the non-tool condition. Object ROIs in LOC were defined as those voxels that responded more to objects (tool + non-tool) than scrambled. Selectivity was measured for each voxel in an ROI by extracting standardized parameter estimates for each condition (relative to fixation).

### Task-dependent functional connectivity

We conducted psychophysiological interaction (PPI; Friston et al., 1997) analyses to examine whether there is task-dependent functional connectivity between dorsal regions involved in computing part relations, and ventral regions involved in object recognition (Friston et al., 1997). A contrastive psychological task covariate was created from the part relations localizer by assigning timepoints corresponding to part-relations blocks a value of 1 and assigning timepoints corresponding to feature blocks a value of -1, then convolving the covariate with a standard HRF. Physiological covariates were generated from each participant’s cleaned residual timeseries by extracting the timeseries from a 6 mm sphere centered on the peak voxel in dorsal ROIs that respond more to the relations than feature condition in the part relations localizer. Finally, a psychophysiological interaction covariate was created for each participant by multiplying the psychological and physiological covariates.

For each participant, 4 runs (randomly selected) of the part relations localizer were used to identify the peak voxel that responded more to the part-relations than feature condition in pIPS and Aips parcels. The cleaned residual timeseries from the left-out two runs were extracted then normalized, concatenated, and then further regressed on the psychological and physiological covariates generated for those runs. A seed-to-whole-brain functional connectivity map was generated by correlating the residual timeseries of every voxel with the interaction covariate, and applying a fisher transform on the resulting map.

Data were analyzed in a cross-validated manner, such that every possible permutation of localizer (4 runs) and left-out runs (2 runs) was used to define the seed region separately, and then analyze connectivity. An average map was created by computing the mean across all permutations and a final group map was created by computing the mean across subjects. Significant voxels were determined by standardizing the group map and applying FDR-correction (*p* < 0.05). Together, this procedure ensures that any correlation between regions is driven by the task-dependent neural interaction, and not by the baseline correlation between regions or shared task activation.

### Effective connectivity analyses

We conducted hypothesis-driven Granger causality analyses (Roebroeck et al., 2005; Seth et al., 2015) to examine the directionality of dorsal and ventral functional connectivity, namely whether the responses in dorsal regions predict those of LOC. The premise underlying Granger causality analyses is as follows. Dorsal cortex will be said to predict the response of ventral cortex if incorporating past responses of dorsal cortex (i.e., *t*-1) improves the prediction of current responses of ventral cortex over above ventral’s own past responses.

Although the low temporal resolution of fMRI precludes strong conclusions about directionality, simulation studies have shown that temporal delays as low as tens of milliseconds can be resolved from the hemodynamic response using Granger causality analyses (Deshpande et al., 2010; Katwal et al., 2009). Thus, by describing the temporal order of events we may gain insight regarding the directionality of information flow between dorsal and ventral cortices.

Cleaned residual timeseries were extracted from a 6 mm sphere centered on the peak voxel in dorsal ROIs that responded more to the relations than feature condition in the part relations localizer. We measured effective connectivity in a task-dependent manner by conducting Granger causality analyses separately on the timeseries from the relations and feature blocks of the part relations localizer.

For each participant, 4 runs (randomly selected) of the part relations localizer were used to identify the peak voxel that responded more to the relations than feature condition in pIPS and aIPS parcels. The cleaned residual timeseries from the left-out two runs were extracted separately from relation and feature blocks, and then concatenated. A single null value was inserted between every block’s timeseries to prevent prediction of temporally discontinuous timepoints. For each dorsal seed region, Granger causality analyses were conducted twice, once with dorsal cortex as the predictor and once with ventral cortex, namely LOC, as the predictor. Following prior work (e.g., Roebroeck et al., 2005), effective connectivity between the areas was calculated by subtracting the dorsal → ventral *F* statistic from the ventral *→* dorsal *F* statistic. A 1-timepoint (i.e., 1 TR) lag was used in all analyses.

Data were analyzed in a cross-validated manner, such that every possible permutation of localizer (4 runs) and left-out runs (2 runs) was used to define the seed region separately, and then analyze connectivity. An average statistic was created by computing the mean *F-*difference for each participant across all permutations. Following previous work, a group analyses were conducted using a Wilcoxon signed-rank test comparing *F*-difference values to 0.

### Experiment 2: Basic-level object categorization in parietal ROIs

We tested whether the multivariate pattern in parietal ROIs that represent object-centered part relations can support basic-level object categorization. We further used representational similarity analyses (RSA), to examine the visual contributions of these ROIs to object recognition. Finally, we used multivariate functional and effective connectivity analyses to examine the degree to which the part relation ROIs in the dorsal pathway interact with the ventral pathway and the degree to which dorsal object responses predict those in ventral cortex.

To this end, participants completed 8 runs (330 s each) during which they viewed images of common objects. The object set was comprised of five categories (boat, camera, car, guitar, lamp) each with five exemplars. Objects were selected from the ShapeNet 3D model dataset (Chang et al., 2015) and rendered to have the same orientation, texture, and color. The original texture and color information was removed to ensure that similarity among objects was on the basis of shape similarity, rather than other features. All stimuli subtended ∼6° visual angle on screen (see Figure 2). To maintain attention, participants performed an orthogonal target detection task wherein they were required to press a button anytime a red box appeared around the object.

**Figure 2.**
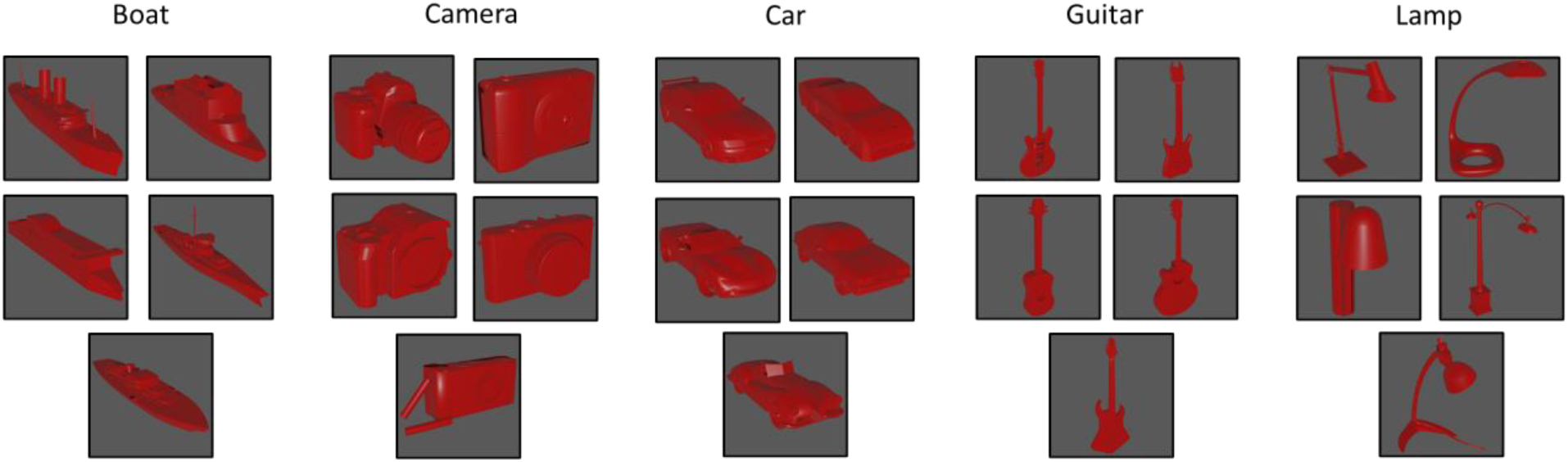
Object stimuli presented in Experiment 2. Participants viewed five exemplars from five categories in an event-related design.

Objects were presented in an event-related design with the trial order and ISI optimized to maximize efficiency using Optseq2 (https://surfer.nmr.mgh.harvard.edu/optseq/). Each stimulus was presented for 1 s, with a jittered ISI between 1 and 8 seconds. Participants viewed 4 repetitions of each object per run. For each participant, parameter estimates for each object (relative to fixation) were extracted for each voxel. Responses to the stimuli in each voxel were then normalized by subtracting the mean response across all stimuli.

#### Representational similarity analyses

A 25 × 25 symmetric neural representational dissimilarity matrix (RDM) was created for each ROI and participant by correlating (1-Pearson correlation) the voxel-wise responses for each stimulus with every other stimulus in a pairwise fashion. Neural RDMs were then Fisher transformed and averaged across participants separately for each ROI. Only the upper triangle of the resulting matrix (excluding the diagonal) was used in subsequent analyses.

Neural RDMs were compared to RDMs created from a model that approximates the spatial relations among component parts, namely a model based on the medial axis shape skeleton. Shape skeletons provide a quantitative description of the spatial arrangement of component parts via internal symmetry axes (Blum, 1973), and are tolerant to variations in the parts themselves (Ayzenberg et al., 2019; Feldman & Singh, 2006). Accumulating research has shown that humans representations of global form are well described by a skeletal model (Lowet et al., 2018), explaining more variance in human responses than conventional ANNs (Ayzenberg et al., 2021; Ayzenberg & Lourenco, 2019) and other descriptors of shape, such as the principal axis (Ayzenberg et al., 2019; Firestone & Scholl, 2014). For our skeletal model, we used a flux-based medial axis algorithm (Dimitrov et al., 2003; Rezanejad & Siddiqi, 2013) which computes a ‘pruned’ skeletal structure tolerant to local variations (Feldman & Singh, 2006). Skeletal similarity between objects was computed as the mean Euclidean distance between each point on one object’s skeleton structure with the closest point on a second object’s skeleton structure.

We also compared neural RDMs for models of low-and high-level vision, namely the Gabor-jet model, a model of image-similarity that approximates the response profile of early visual regions (Margalit et al., 2016), and the penultimate layer of CorNet-S, a recurrent artificial neural network designed to approximate the response profile of the ventral visual pathway in monkeys (Kubilius et al., 2019). Object similarity for both Gabor-jet and CorNet-S were computed as the mean Euclidean distance between feature vectors for each object image (see Figure 9).

#### Multivariate connectivity analyses

We conducted multivariate pattern dependence (MVPD) analyses (Anzellotti et al., 2017) to examine whether dorsal ROIs involved in computing part relations interact with ventral object regions during object viewing. MVPD tests the degree to which the multivariate activation timeseries of a seed region accounts for the variance of the multivariate activation timeseries of a target region.

For each participant, data were split into a training (6 runs) and test (2 runs) set. A multivariate timeseries was generated from each participant’s cleaned residual timeseries training data by extracting the timeseries of each voxel from a 6 mm sphere centered on the peak voxel in dorsal ROIs that responds more to the part-relations than feature blocks in the object-centered relations localizer. The dimensionality of the voxel timeseries was then reduced by applying principal components analysis (PCA) and selecting the components that explain 90% of the variance. The same procedure was then repeated for a target region using a searchlight with 6 mm sphere. Next, using the training data, a linear regression was fit separately on each component of the target region using the components from the seed region as predictors. This procedure results in a series of beta weights describing the linear mapping between the principal components of the seed region to each individual principal component of the target region. For computational efficiency, the searchlight was conducted within an extended visual cortex mask created using an atlas from Wang et al. (2014) comprised of occipital, dorsal, and ventral visual cortices.

The beta weights from the training data are then used to generate a predicted multivariate timeseries for left-out runs of the target region, which is then correlated (Pearson) with the actual observed timeseries of the target region. A final fit value is computed as the weighted mean of correlations across target region principal components, with the weighting of each correlation determined by the proportion of variance explained by each target component. A single map for each participant is created by averaging the weighted correlations following 5-fold cross-validation, and then Fisher transforming the correlations. A final group map is created by computing mean across participants. Significant voxels were determined by standardizing the group map and applying FDR-correction (*p* < 0.05).

#### Multivariate effective connectivity

We conducted hypothesis-driven multivariate Granger causality analyses to examine the directionality of functional connectivity between dorsal and ventral pathways. Like its univariate counterpart, multivariate Granger causality tests whether past responses of one multivariate timeseries (e.g., dorsal cortex) predict the current responses of a second multivariate timeseries (e.g., LOC) over and above their own past timepoints.

For each participant, the entire cleaned residual timeseries (8 runs) was extracted from a 6 mm sphere centered on the peak voxel in dorsal ROIs that responds more to the part-relations than feature blocks in the object-centered relations localizer. The dimensionality of the voxel timeseries was then reduced by applying principal components analysis (PCA) and selecting the components that explain 90% of the variance. The same procedure was then repeated for LOC. To conduct multivariate Granger causality, the total number of components for each ROI was matched to the ROI with fewer components.

For each dorsal seed region, multivariate Granger causality was conducted twice, once with dorsal seed region as the predictor and once with LOC as the predictor. As in univariate Granger causality, effective connectivity between the two regions was calculated by subtracting the dorsal → ventral *F* statistic from the ventral *→* dorsal *F* statistic. A 1-timepoint (i.e., 1 TR) lag was used in all analyses. Group analyses were conducted using a Wilcoxon signed-rank test comparing *F*-difference values to 0.

## Results

### Experiment 1: Selectivity for object-centered relations in the dorsal pathway

#### ROI definition

See Table 1 for a summary of significant group-level clusters from every localizer. The part relations localizer (4 runs) identified significant clusters in pIPS and aIPS in the right hemisphere (rpIPS, raIPS) of every participant and in 10 out of 12 participants in the left hemisphere (lpIPS, laIPS; see Figure 3A). Likewise, a group averaged map created using 2 runs (left out to measure selectivity) from every participant also revealed significant clusters in pIPS and aIPS, though these were found exclusively in the right hemisphere (see Figure 3B).

**Table 1.** Significant group level clusters for the object-centered part relations, allocentric spatial relations, and tool localizer. MNI Coordinates correspond to the peak voxel within each cluster. The depth localizer is not listed because there were no significant clusters at the group level.

| Localizer | Region | MNI Coordinate |  |  |
| --- | --- | --- | --- | --- |
|  |  | x | y | z |
| Object part relations |  |  |  |  |
|  | 1 R Posterior Intraparietal Sulcus (IPS0) | 26 | -76 | 44 |
|  | 2 R Ventral Intraparietal Complex (VIP) | 22 | -58 | 64 |
|  | 3 R Middle Temporal Area (MT) | 42 | -78 | 12 |
|  | 4 R Temporal Parietal Junction (TPJ) | 52 | -60 | -2 |
| Allocentric spatial relations |  |  |  |  |
|  | 1 L Intraparietal sulcus (IPS1) | -26 | -72 | 24 |
|  | 2 L Ventral Intraparietal Complex (VIP) | -16 | -68 | 58 |
|  | 3 R Ventral Intraparietal Complex (VIP) | 16 | -62 | 56 |
|  | 4 L Secondary Somatosensory Cortex (S2) | -38 | -38 | 48 |
|  | 5 R Secondary Somatosensory Cortex (S2) | 45 | -40 | 63 |
|  | 6 R V3A/V3B | 34 | -78 | 16 |
|  | 7 L Middle Temporal Area (MT) | -48 | -72 | 2 |
|  | 8 L Fundal Superior Temporal (FST) | -48 | -66 | -6 |
| Tools |  |  |  |  |
|  | 1 L Lateral Interparietal Area (LIP) | 24 | -58 | 64 |
|  | 2 R Ventral Intraparietal Complex (VIP) | -22 | -54 | 58 |
|  | 3 L Middle Temporal Area (MT) | -46 | -76 | 6 |
|  | 4 L Temporal Parietal Junction (TPJ) | -58 | -72 | 0 |
|  | 5 R Temporal Parietal Junction (TPJ) | 56 | -68 | 4 |

**Figure 3.**
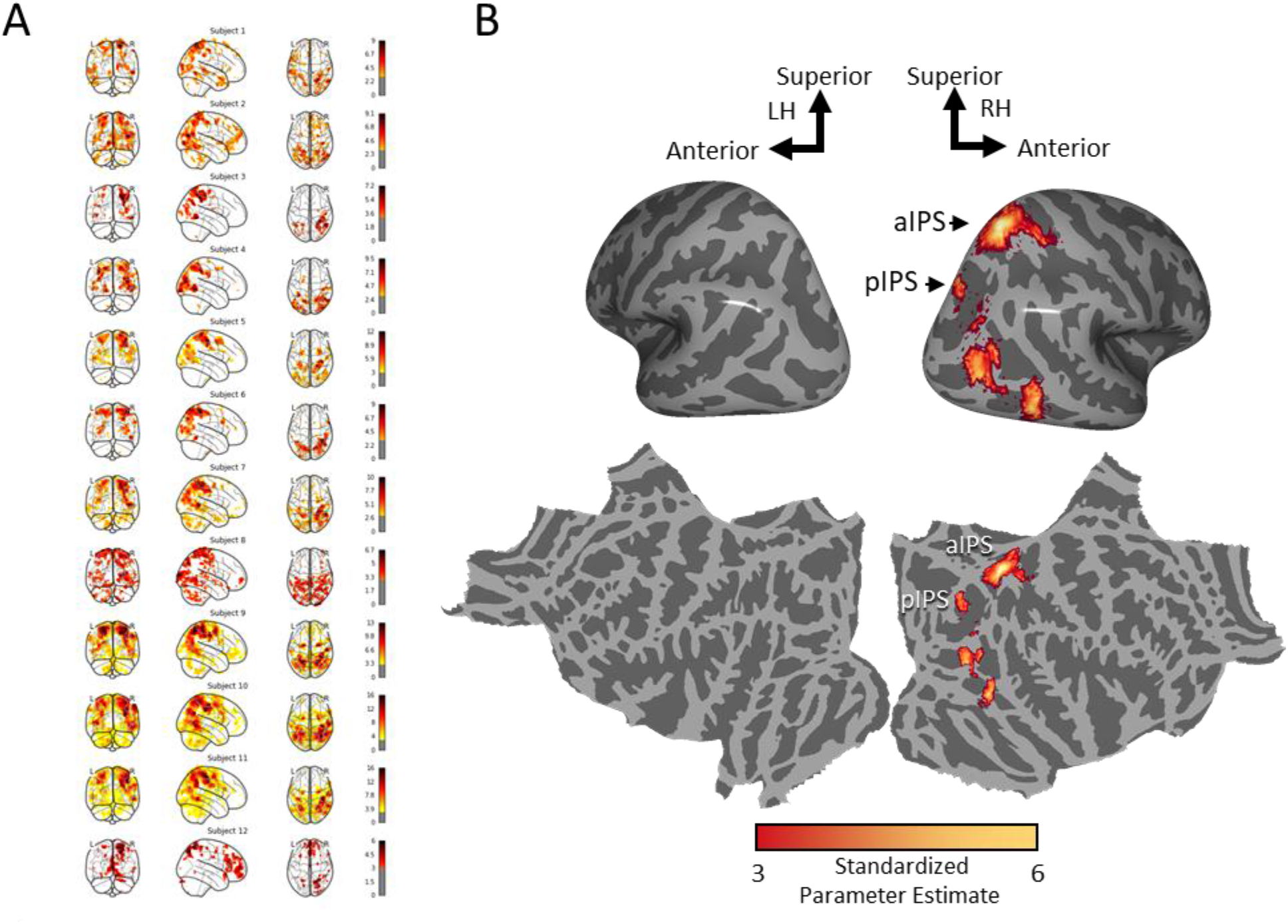
Significant activation to part relations (versus features) condition from the object-centered part relations localizer displayed (A) for each individual participant and in (B) a group average map inflated (above) and flattened (below). Values reflect the standardized parameter estimate.

#### Selectivity for part relations

To test whether these ROIs are *selective* for object-centered part relations, we examined the response in this region (relative to fixation; see Material and Methods) to (1) activation in the relations blocks of the part relations localizer (independent runs), as well as the other dorsal conditions, namely, (2) distance as determined from the allocentric relations localizer, (3) 3D shape from the depth localizer, and (4) tools from the tool localizer.

A repeated-measures ANOVA with ROI (pIPS, aIPS), hemisphere (left, right), and condition (part relations, distance, 3D shape, tools) as within-subjects factors revealed that there was a significant main-effect of condition, *F*(3, 24) = 8.26, *p* < .001, η_p_2 = 0.53. There were no other main-effects or interactions (*ps* > .102). Post-hoc comparisons (Holm-Bonferroni corrected) revealed that activation to the part-relations condition was higher than distance (*t*[11] = 4.64, *p* < .001, *d* = 1.55), 3D shape (*t*[11] = 4.16, *p* = .002, *d* = 1.39), and tool (*t*[11] = 4.48, *p* = .008, *d* = 1.16) conditions.

Thus, these analyses suggest that the dorsal pathway represents object-centered part relations, and that this representation is independent of allocentric spatial relations and other object properties represented by the dorsal pathway.

Although these analyses did not reveal a significant difference between left and right hemisphere ROIs, examination of the group map suggests that the part relations may be more strongly represented in the right hemisphere. To explore these possible differences, we also analyzed each ROI separately. Note, due to the exploratory nature of this analysis, these results should be interpreted with caution.

Separate repeated measures ANOVAs were conducted for participants’ left and right pIPS and aIPS which revealed main-effects of condition in all four regions (lpIPS: *F*[3, 33] = 3.92, *p* = .021, η_p_^2^ = 0.33; rpIPS: *F*[3, 33] = 8.70, *p* < .001, η_p_^2^ = 0.44; laIPS: *F*[3, 33] = 4.69, *p* = .009, η_p_^2^ = 0.34; raIPS: *F*[3, 33] = 12.57, *p* < .001, η_p_^2^ = 0.53), with the response to part relations numerically highest in each region (see Figure 4). However, post-hoc comparisons (Holms-Bonferroni corrected) revealed that activation to part relations was statistically highest only in the right hemisphere parietal regions, but not the left hemisphere parietal regions. Namely, in the right hemisphere, the activation to part relations was significantly higher than distance (rpIPS: *t*[11] = 4.66, *p* < .001, *d* = 1.34; raIPS: *t*[11] = 4.18, *p* < .001, *d* = 1.21), 3D shape (rIPS: *t*[11] = 3.47, *p* = .006, *d* = 1.00; raIPS: *t*[11] = 5.77, *p* < .001, *d* = 1.67), and tools (rpIPS: *t*[11] = 4.05, *p* = .001, *d* = 1.17; raIPS: *t*[11] = 4.52, *p* < .001, *d* = 1.31). By contrast, in the left hemisphere, pIPS responses to part relations were higher than distance (*t*[11] = 3.21, *p* = .023, *d* = 1.07), but not 3D shape or tools (*ts* < 2.65, *ps >* .071, *ds* < 0.88). In left aIPS, responses were higher than distance (*t*[11] = 3.51, *p* = .010, *d* = 1.1) and 3D shape (*t*[11] = 2.87, *p* = .039, *d* = 0.91), but not tools (*t*[11] = 2.39, *p* = .097, *d* = 0.75). In combination with the group statistical map (Figure 3), these results suggest that object-centered part relations may be represented more strongly in the right than left hemisphere parietal regions.

**Figure 4.**
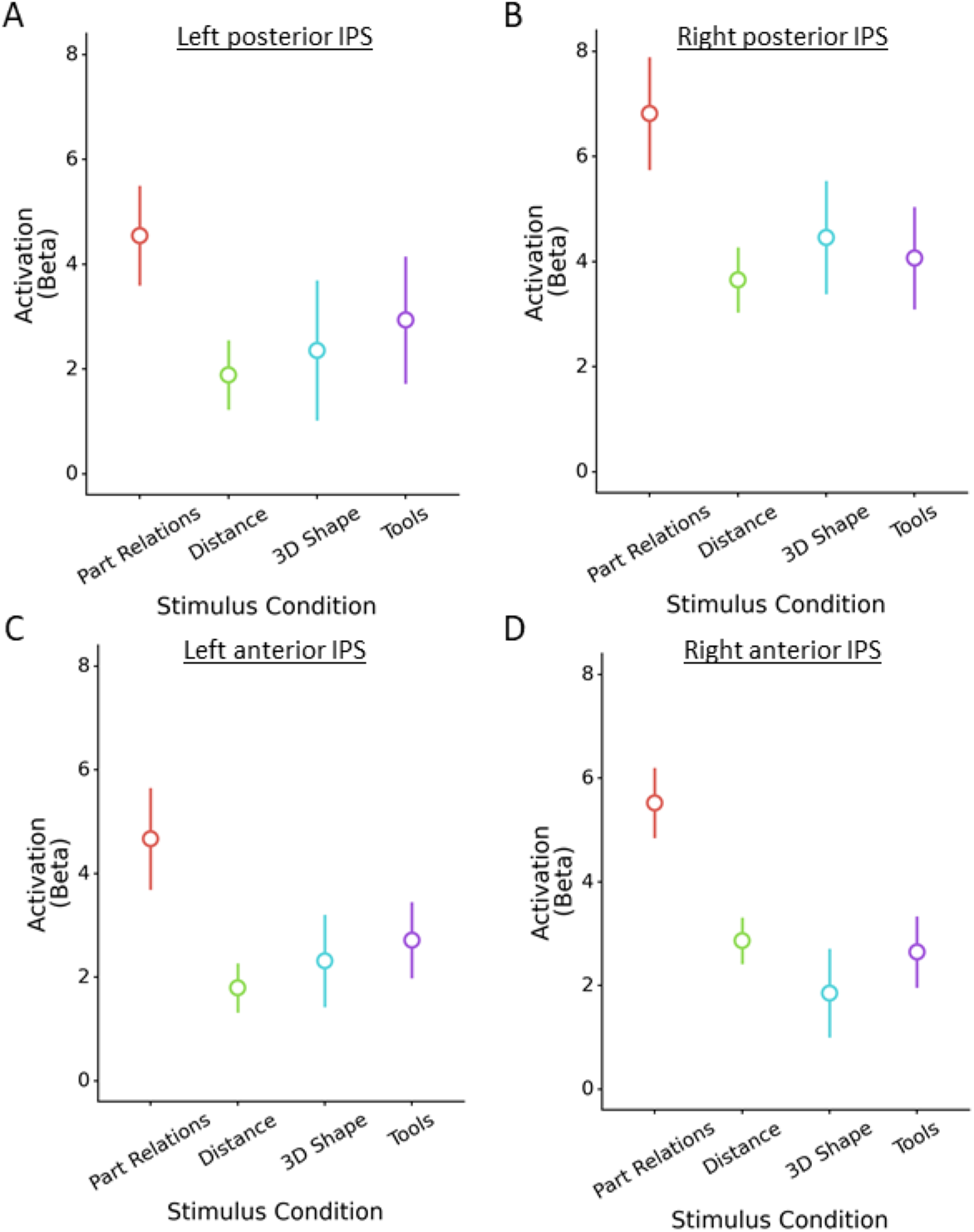
Activation to the part relations (left-out runs), allocentric distance, 3D shape, and tools conditions in (A) left pIPS and (B) right pIPS, (C) left aIPS, and (D) right aIPS. Activation values reflect the standardized parameter estimate. Error bars reflect standard error of the mean.

#### Conjunction analyses

To explore further the degree to which parietal regions involved in computing part relations overlap with regions computing other dorsal properties, we conducted whole-brain conjunction analyses. First, group-averaged statistical maps were created for every localizer and a cluster-correction threshold applied (*p* < .001; see Table 1). The resulting statistical maps were consistent with prior research on the neural basis of the allocentric relations (Zachariou et al., 2017) and of tool representations (Chen et al., 2016; Gallivan et al., 2013). No significant clusters were found for the activation profiles on the depth localizer (Georgieva et al., 2008).

Next, we calculated the proportion of independent and overlapping voxels by converting the thresholded statistical map from each group-averaged localizer into binary masks and overlaying them with the thresholded statistical map from part relations localizer. Binomial tests revealed that, in right pIPS, there were significantly more independent than overlapping voxels that responded to part relations. Here, the allocentric relations ROI had the greatest amount of overlap with part relations ROI in pIPS (overlapping voxels: 42%, *p* < .001). There were no overlapping voxels from the depth or tool ROIs above the cluster corrected threshold. By contrast, in right aIPS, there were significantly more voxels that overlapped with the allocentric relations ROI than were independent (overlapping voxels: 65%, *p* < .001). There was also overlap with the tool ROIs (overlapping voxels: 43%, *p* < .001), but there were significantly more independent voxels than overlapping ones. There were no overlapping voxels with the depth localizer (0%). Together these results suggest part relations may be represented along a gradient within the dorsal pathway, with both distinct and overlapping components.

Finally, to visualize this gradient better, statistical maps were converted into proportions, such that, for each voxel, a value closer to 1 indicates a greater response to part relations and a value closer 0 indicates a greater response to one of the other dorsal properties (e.g., allocentric relations; see Figure 5). Consistent with the analyses above, these maps reveal the least overlap between part relations and other dorsal ROIs in pIPS and the most overlap in aIPS.

**Figure 5.**
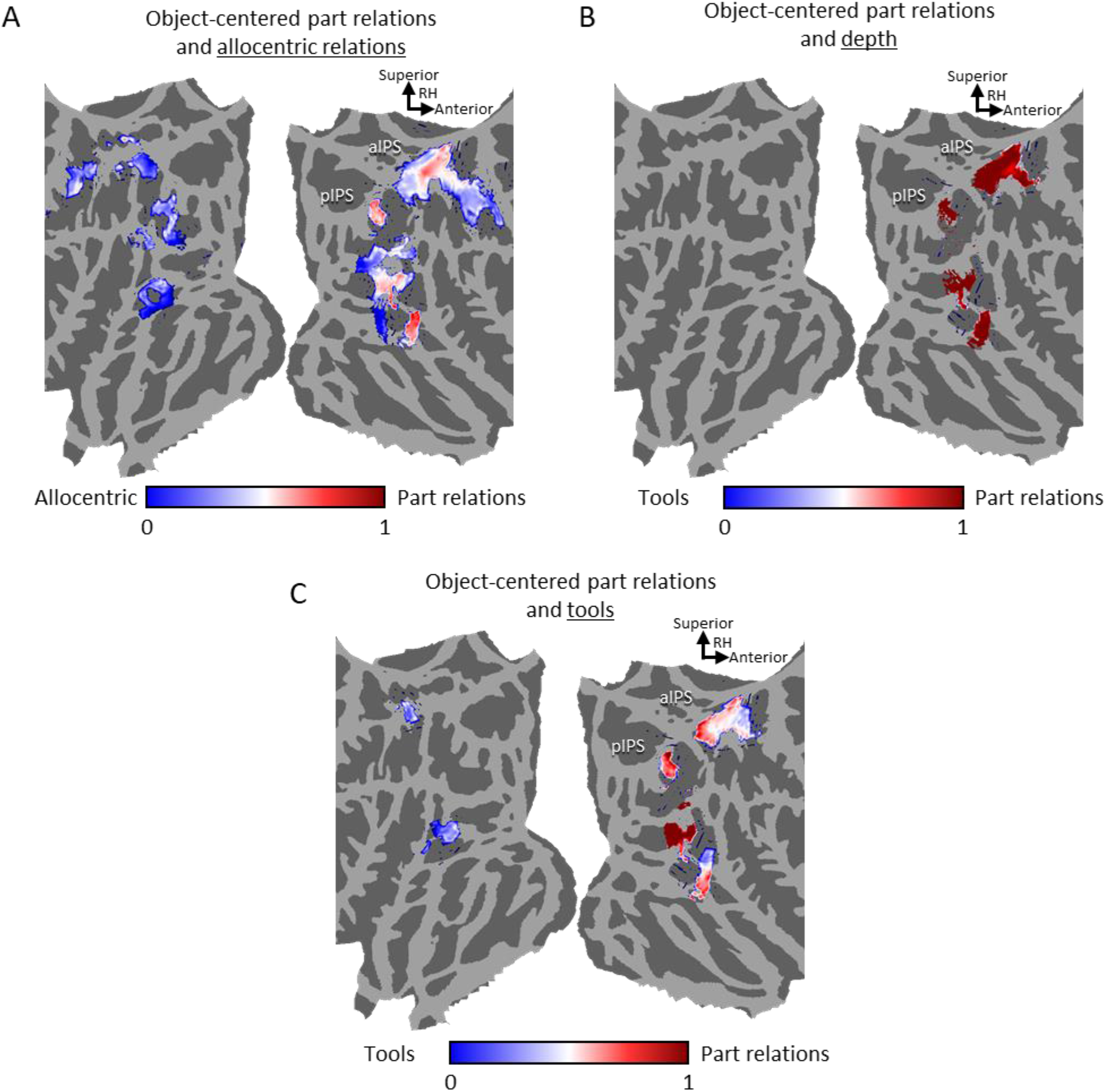
Conjunction maps illustrating areas of distinct and overlapping coding for object-centered part relations and (A) allocentric relations, (B) depth, and (C) tools. A value closer 1 indicates a greater response to part relations; a value closer to 0 indicates a greater response to the control localizer. Maps are zoomed in on the visual cortex for easier inspection.

#### Task-dependent functional connectivity

If the role of the dorsal pathway in object recognition is to compute object-centered part relations, then a prediction is that pIPS and aIPS will also be functionally connected to the ventral pathway – the nexus of object recognition processing. More specifically, the prediction is that functional connectivity between right and left pIPS or aIPS with ventral cortex will depend on the task demands, such that connectivity would be greatest when perception of part relations is needed, as in the relations, but not feature, condition of the localizer. To test this prediction, we conducted PPI analyses to examine whether there was task-dependent functional connectivity between left and right pIPS and aIPS regions involved in computing object-centered part relations, and ventral regions involved in object recognition (see Materials and Methods).

Examination of the group map (Figure 6) revealed significant connectivity between right hemisphere pIPS and aIPS with bilateral ventral pathway regions. Interestingly, there was relatively little connectivity with other dorsal regions, suggesting that the function of right hemisphere pIPS and aIPS may be specifically in the service of object recognition processes in the ventral pathway rather than action processes in other dorsal regions. There was no significant connectivity with left pIPS or aIPS that survived FDR correction.

**Figure 6.**
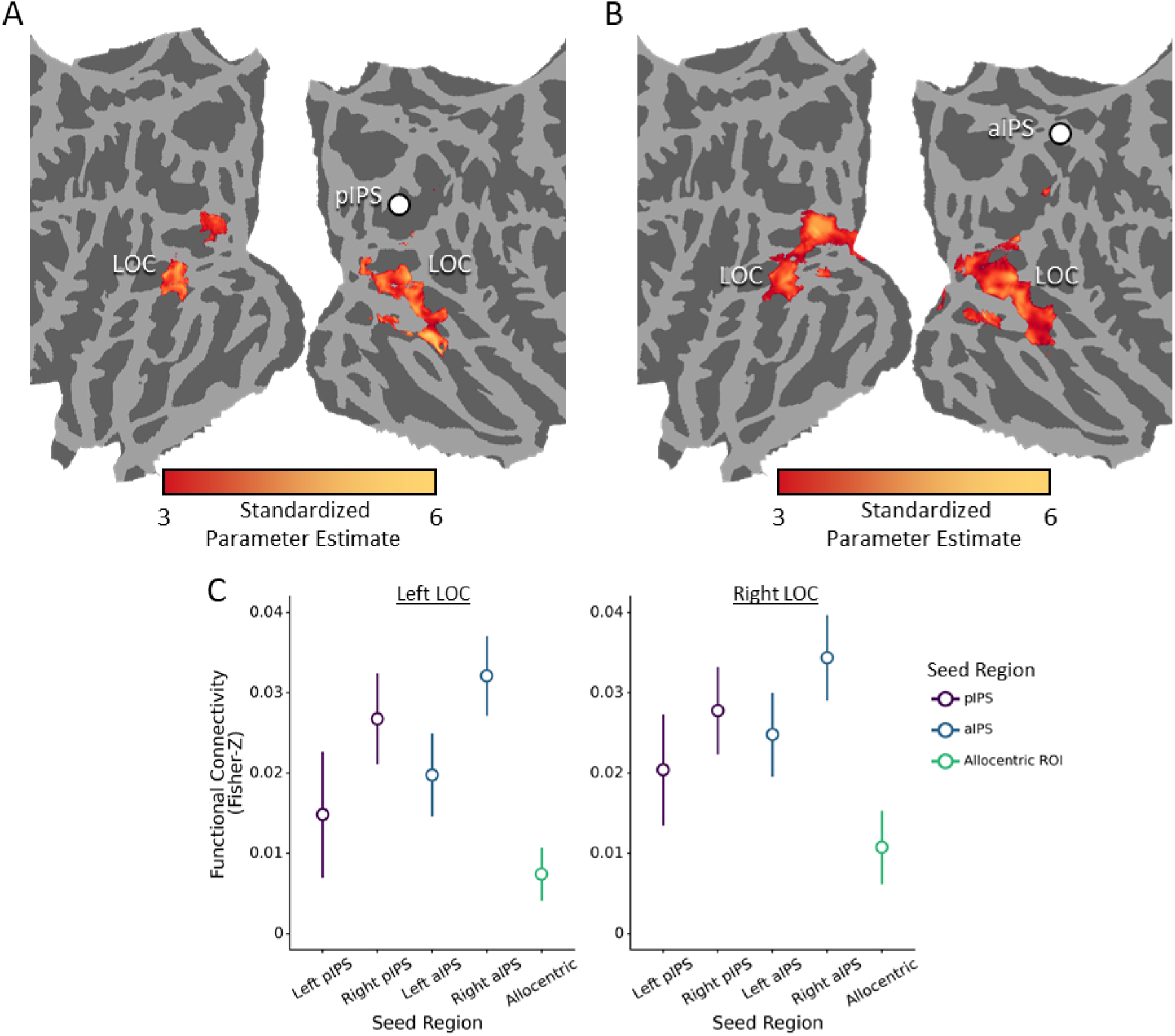
Task-based functional connectivity results. (A-B) Functional connectivity map (zoomed in on the visual cortex) for (A) right pIPS and (B) right aIPS. Seed regions are displayed as white circles. There was no functional connectivity above the cluster corrected threshold in left pIPS, left aIPS, or the left allocentric ROI. (C) Plots comparing the connectivity between pIPS, aIPS, and the other ROIs in left LOC and right LOC ROIs. Error bars reflect standard error of the mean.

To further examine the specificity of task-dependent connectivity to these regions, we reanalyzed the data from the part relations localizer using the peak voxel from the allocentric relations ROI in the left hemisphere as our seed region. This ROI was chosen because it does not overlap with part relations ROIs, but nevertheless has a conceptually similar representation. These analyses revealed no significant connectivity between allocentric relations ROIs in the left hemisphere and the ventral visual pathway. Moreover, a direct comparison between regions (Holm-Bonferroni corrected), revealed that task-dependent connectivity with LOC, a ventral object region, was significantly stronger with right pIPS (lLOC: *t*(11) = 3.41, *p* = .005, *d* =0.99; rLOC: *t*(11) = 3.28, *p* = .007, *d* =0.95) and aIPS (lLOC: *t*(11) = 4.36, *p* < .001, *d* =1.26; rLOC: *t*(11) = 4.56, *p* < .001, *d* =1.32) than left allocentric relations ROIs. There were no differences in connectivity between the other pIPS and aIPS regions (*ps* > .217). Together, these findings suggest that dorsal regions involved in computing object-centered part relations, particularly in the right hemisphere are preferentially connected to the ventral stream to support object recognition.

#### Task-dependent effective connectivity

If dorsal regions propagate information about object-centered part relations to the ventral pathway for recognition, then one should expect that representations of part relations in pIPS and aIPS will temporally precede and will predict those in ventral cortex. More specifically, the prediction is that the past timepoints of pIPS or aIPS will predict current timepoints of ventral cortex over and above ventral’s own past time points. Moreover, this effect should be strongest for the relations condition of the localizer, not the feature condition. To test this prediction, we conducted Granger causality analyses to examine the effective connectivity between left and right pIPS and aIPS regions involved in computing object-centered part relations and LOC involved in object recognition (see Materials and Methods).

A Wilcoxon signed-rank comparison to 0 revealed significant effective connectivity during the relations blocks between left pIPS with right LOC (*W* = 74, *p* = .002, *d* = 0.90), but not left LOC (*W* = 57, *p* = .088, *d* = 0.46), and between right pIPS with right LOC (*W* = 66, *p* = .017, *d* = 0.70), but not left LOC (*W* = 45, *p* = .339, *d* = 0.15) (see Figure 7). There was positive effective connectivity between right aIPS with left (*W* = 60, *p* = .055, *d* = 0.54) and right (*W* = 59, *p* = .065, *d* = 0.51) LOC during the relations blocks, although these effect did not reach the criteria for significance. There were no significant effects for left aIPS for the relations blocks in either left or right LOC (*Ws* < 46, *ps* > .311, *ds* < 0.18), nor any of the ROIs in the feature blocks (*Ws* < 56, *ps* > .102, *ds* < 0.44).

**Figure 7.**
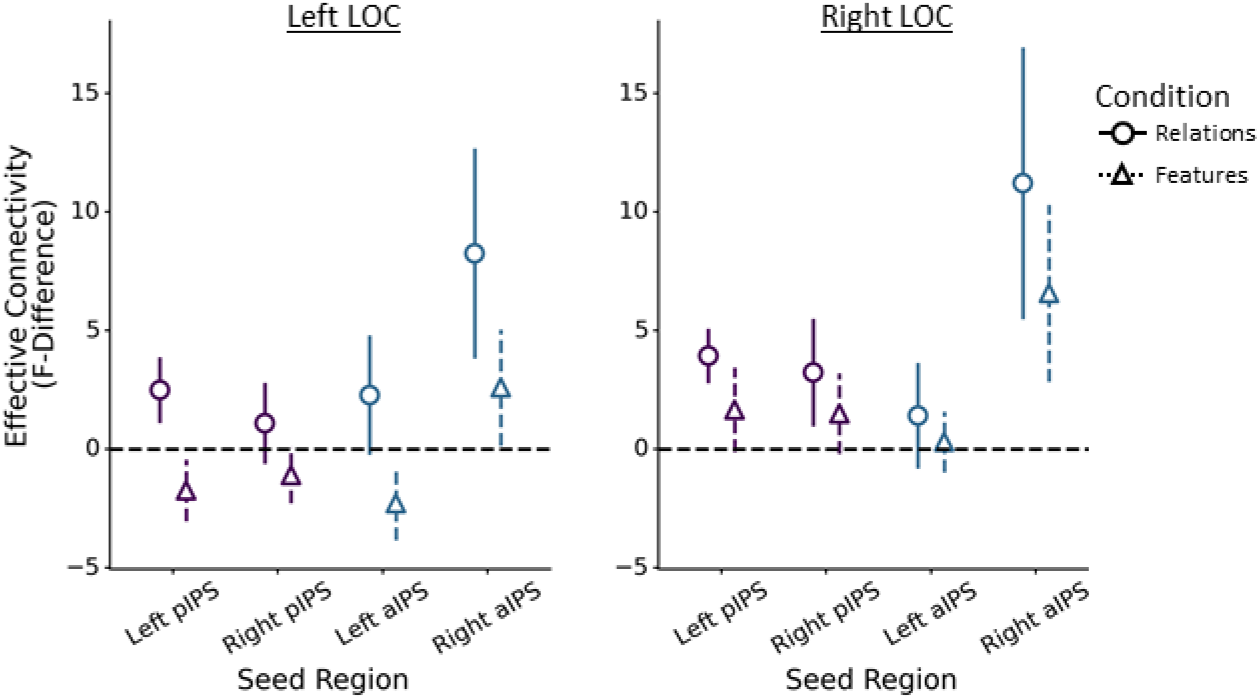
Plots comparing the task-based effective connectivity between left and right pIPS and aIPS with left LOC and right LOC ROIs. Error bars reflect standard error of the mean.

Separate repeated-measures ANOVAs were further conducted to analyze effective connectivity as a function of ROI (pIPS, aIPS), hemisphere (left, right), and condition (relations, features). As hypothesized, these analyses revealed a significant main-effect of condition, such that effective connectivity was overall higher for the relations than feature blocks in left LOC, *F*(1, 11) =7.45, *p* = .020, η_p_^2^ = 0.40, though right LOC did not meet criteria for significance, *F*(1, 11) =3.60, *p* = .084, η_p_^2^ = 0.25. Moreover, there was a significant ROI × hemisphere interaction in both left LOC, *F*(1, 11) =5.46, *p* = .039, η_p_^2^ = 0.33, and right LOC, *F*(1, 11) =7.26, *p* = .019, η_p_^2^ = 0.41, such that effective connectivity was higher in right aIPS than left aIPS. However, none of the post-hoc comparisons were significant following Holm-Bonferroni correction (*ps* > .066). Together, these findings suggest that pIPS and aIPS transmit information about object-centered part relations to the ventral pathway, rather than the other way around.

#### Analysis on larger sample

All findings from Experiment 1 were replicated successfully with a larger sample (*n* = 18). The part relations localizer (4 runs) identified significant clusters in pIPS and aIPS in all 18 participants in the right hemisphere, but 14 participants exhibited left pIPS ROI and 16 exhibited left aIPS ROI. We found selectivity for object-centered part relations in right pIPS and aIPS, with responses greater than allocentric relations, 3D shape, and tools, (*ps* < 0.006). Moreover, we found significant task-based functional connectivity between right pIPS and aIPS with both left and right LOC, which was greater than a control region defined using allocentric relations (*ps* < .008). Finally, we found significant effective connectivity between right pIPS with right LOC (*p* = .048) during the relations, but not feature blocks of the part-relations localizer. Importantly, there was a main effect of condition in left LOC (*p* = .010), such that there was overall greater effective connectivity during the relations blocks than the feature blocks.

### Experiment 2: Dorsal contributions to object recognition

#### Category decoding

To test whether dorsal regions that compute object-centered part relations contribute to object recognition, we examined whether multivariate pattern within these regions could be used to classify objects (see Figure 2). Using a 20-fold cross-validation procedure, a Support Vector Machine (SVM) classifier was trained on the multivariate pattern for three exemplars from each category, and then tested on the category of the two left out exemplars.

One-sample comparisons to chance (0.20) revealed that category decoding was significantly above chance in right pIPS, *M* = 32.7%, *t*(11) = 3.15, *p* = .009, *d* = 0.91, but not in right aIPS, left pIPS or left aIPS ROIs defined on the basis of part relations (*Ms* < 23.4%, *ps* > .110, *ds* < 0.72; Figure 8). To further examine the specificity of category decoding in dorsal regions, we also tested how well a left hemisphere allocentric relations ROI can decode object categories. These analyses revealed that decoding was not above chance in the left allocentric ROI, *M* = 18.7%, *t*(11) = -0.82, *p* = .780, *d* = 0.23. Direct comparisons between right pIPS and the other regions (Holm-Bonferroni corrected) further confirmed that, categorization accuracy was significantly higher in right pIPS than left allocentric regions (*t*[11] = 3.88, *p* = .004, *d* = 1.23 and left aIPS (*t*[11] = 4.32, *p* = .001, *d* = 1.37), though not right aIPS (*t*[11] = 2.65, *p* = .096, *d* = 0.837) nor left pIPS (*t*[11] = 2.48, *p* = .127, *d* = 0.78). Next, we examined how category decoding in the dorsal pathway compares to ventral pathway object recognition regions, namely LOC. As would be expected, categorization accuracy was above chance in left and right LOC, (lLOC: *M* = 26.0%, *t*[11] = 3.56, *p* = .004, *d* = 1.03; rLOC: *M* = 26.7% *t*[11] = 2.30, *p* = .042, *d* = 0.66), with the neither region differing significantly from right pIPS (*ts* < 1.62, *ps* > .357, *ds* < 0.42). Thus, regions in right posterior IPS involved in computing object-centered part relations can support categorization of object exemplars.

**Figure 8.**
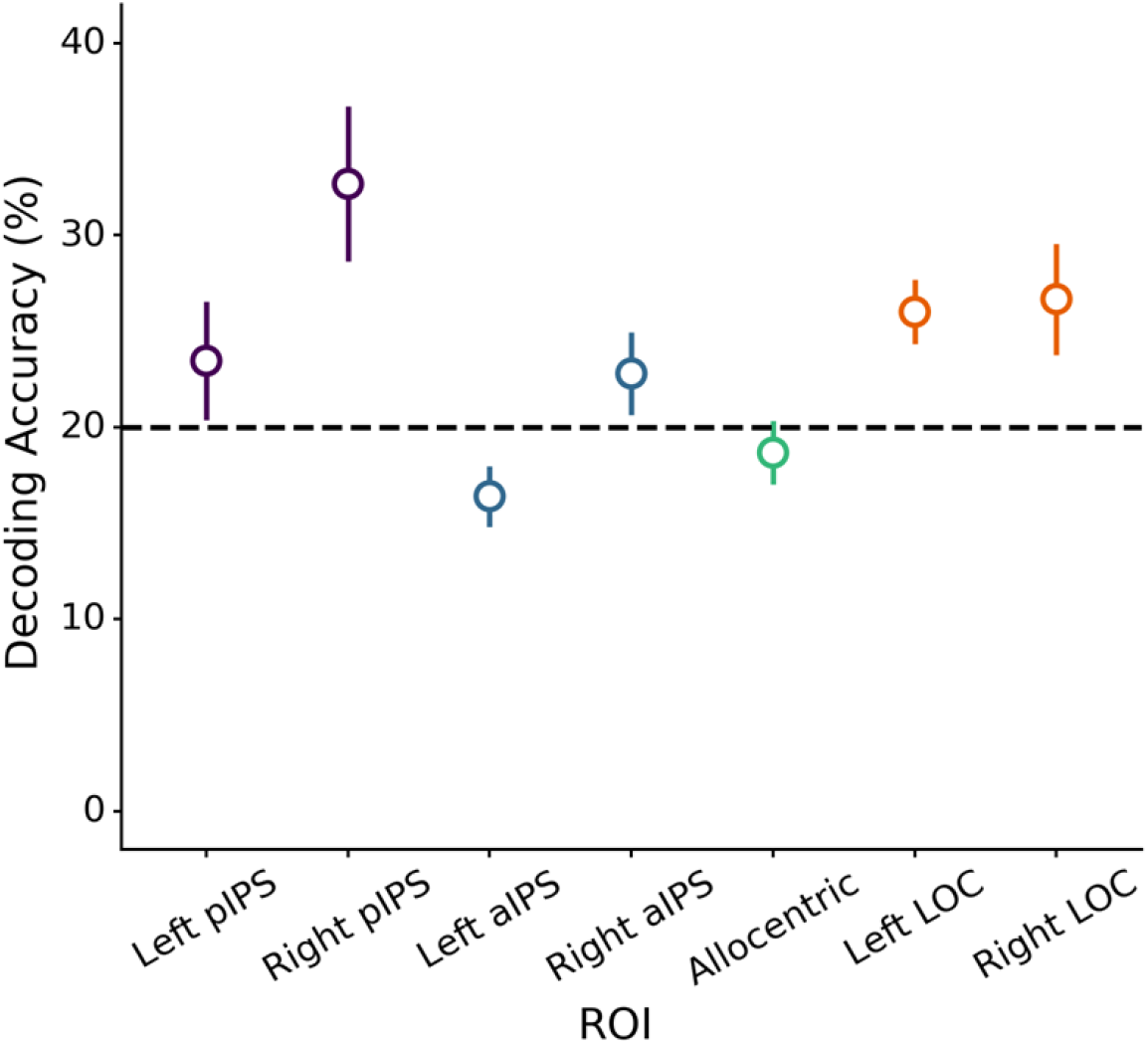
Object categorization accuracy for pIPS, aIPS, the left allocentric ROI, and LOC. Error bars reflect standard error of the mean.

#### Representational content of dorsal ROIs

The results above show that a region in pIPS defined on the basis of part relations can be used to decode the category of objects. Yet, despite the fact that this region was defined using a part relations localizer, it is possible that categorization was accomplished using other visual properties. Indeed, it is well known that pIPS retains a retinotopic organization (Wang et al., 2014) and is tightly connected to early visual cortex (Greenberg et al., 2012). Thus, it is possible that the categorization performance of right pIPS may have been achieved on the basis of low-level image-level similarity. Moreover, it is unclear to what degree categorization in right pIPS is accomplished using high-level visual representations distinct from those in the ventral pathway.

To examine whether right pIPS accomplished object categorization on the basis of object-centered part relations, we used representational similarity analyses (RSA). Specifically, we tested whether a skeletal model, which approximates object-centered part relations, explains unique variance in pIPS over and above other models of vision (see Materials and Methods). Like the representation measured by the part relations localizer, skeletal models describe the spatial arrangement of object parts while ignoring variations in the parts themselves (see Figure 9). Indeed, skeletal models explain more variance in participants judgments of part relations than other models of vision (Ayzenberg & Lourenco, 2019; Lowet et al., 2018).

**Figure 9.**
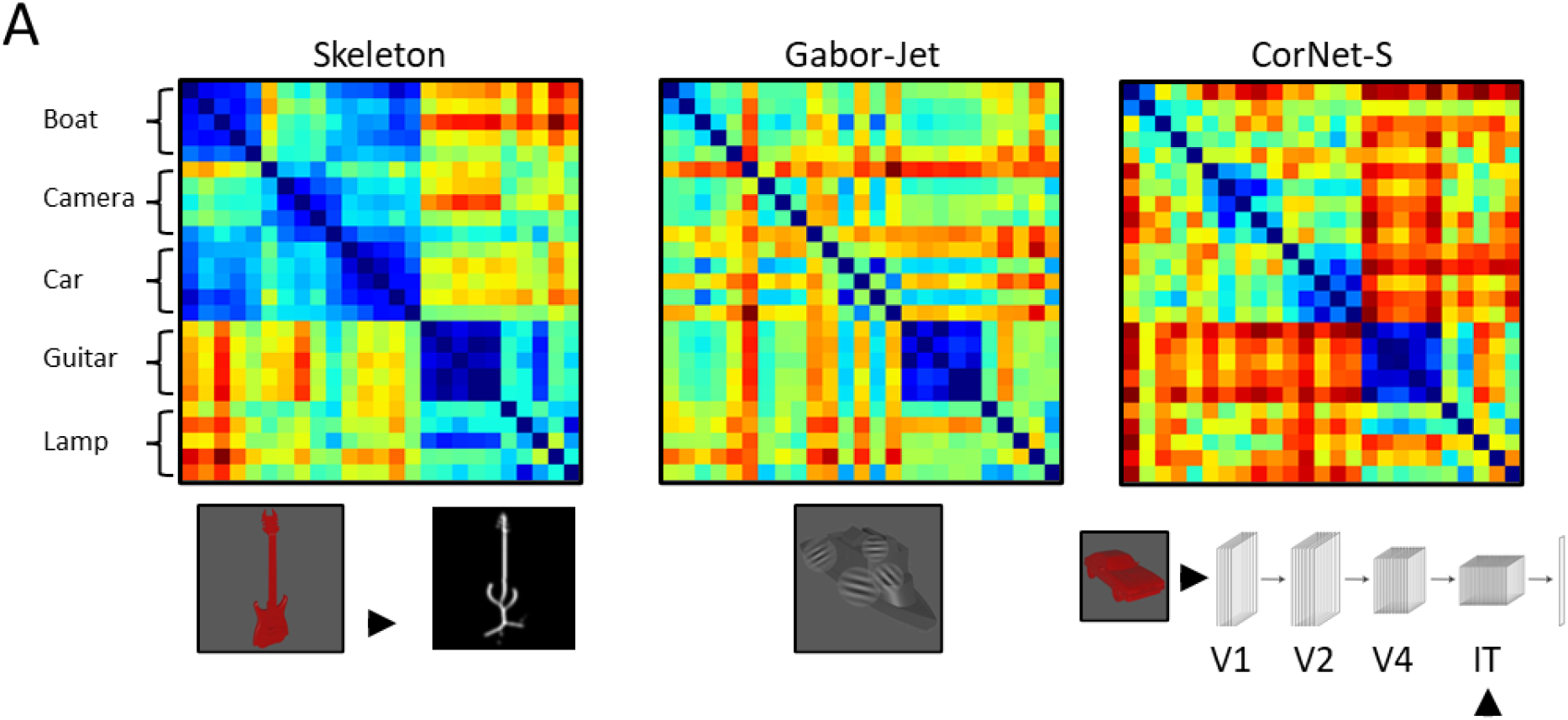
Representational dissimilarity matrices (RDMs) and a schematic illustration of the (left) the skeletal model, (middle) Gabor-jet model, and (right) CorNet-S.

As a comparison, we also tested whether ROIs are well described by Gabor-jet (GBJ), a model of low-level image similarity (Margalit et al., 2016; see Figure 9), as well as CorNet-S a neural network model whose upper layers approximate the response profile of high-level ventral regions in monkeys (Kubilius et al., 2019; Schrimpf et al., 2018; see Figure 9).

To test whether the skeletal model explained unique variance in right pIPS, we conducted linear regression analyses with the neural RDM from pIPS as the dependent variable and the different models of visual similarity as predictors (Skeleton ∪ GBJ ∪ CorNet-S; see Figure 9). Consistent with the localizer results of Experiment 1, these analyses revealed that only skeletal model explained unique variance in right pIPS (*β* = 0.33, *p* < .001), not the other models (GBJ: *β* = 0.04, *p* = .493; CorNet-S: *β* = -0.02, *p* = .839). The skeletal model also explained the most variance in right aIPS, though it approached but did not meet the criteria for statistical significance (skeleton: *β* = 0.14, *p* = .068; GBJ: *β* = 0.00, *p* = .968; CorNet-S: *β* = -0.07, *p* = .376). The skeletal model did not explain significant unique variance in any other dorsal ROI (*β* < 0.12, *ps* > .113; see Figure 10A-B). These findings are consistent with the results of Experiment 1, which suggest that pIPS and aIPS ROIs, particularly those in the right hemisphere, represent objects in terms of their object-centered part relations. Moreover, these results suggest that categorization in right pIPS was accomplished by representing part relations, not other low-or high-level visual properties.

**Figure 10.**
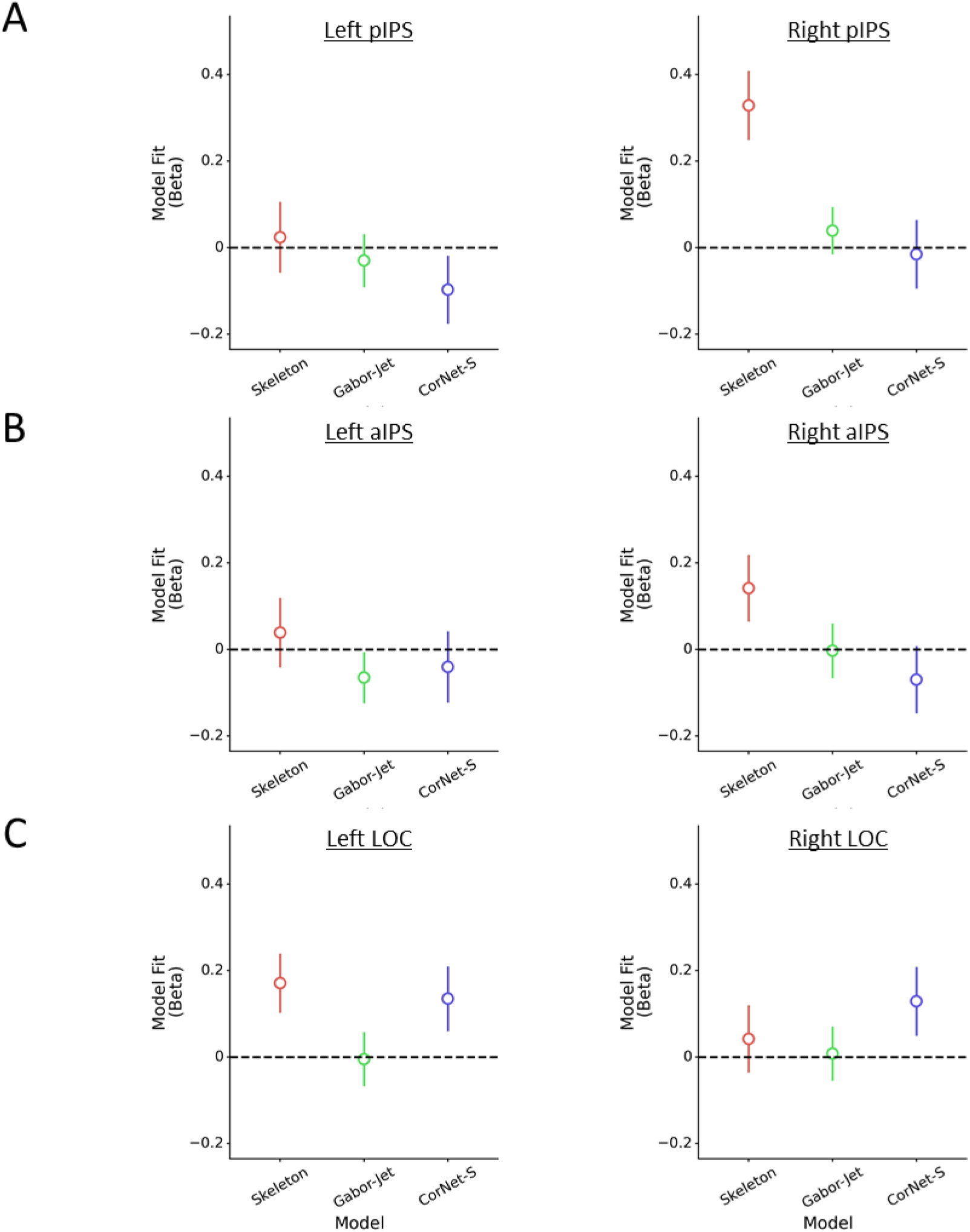
Results of the representational similarity analyses (RSA). (A-C) Standardized coefficients (Betas) from the linear regression analyses examining the fit of the skeletal, Gabor-jet, and CorNet-S models for left and right (A) pIPS, (B) aIPS, and (C) LOC.

#### Unique contributions of dorsal ROIs to ventral processing

Next, we examined whether right pIPS represents distinct visual information from ventral object regions such as LOC. We repeated the linear regression analyses, except here we used neural RDMs from left and right LOC as the dependent variable. These analyses revealed that, the skeletal model explained unique variance in left (*β* = 0.17, *p* = .023), but not right LOC (*β* = 0.04, *p* = .582; see Figure 10C).

Although in Experiment 1 we found that coding of part relations in the dorsal pathway precedes the ventral pathway, this finding nevertheless raises the question: do regions of dorsal cortex compute object-centered part relations and then transmit that information to ventral cortex for object recognition? Or, are part relations computed in the ventral pathway, as previously proposed (Ayzenberg et al., 2021; Behrmann et al., 2006) and transmitted to dorsal regions such as right pIPS? Alternatively, part relations may be coded in parallel in both pathways. To investigate these possibilities, we examined whether multivariate response in pIPS mediates the relation between the skeletal model and the neural RDM in LOC. In other words, we tested whether skeletal coding in LOC is represented independently or by way of right pIPS.

To test these possibilities, we first repeated the linear regression analyses in left LOC, but this time we included the neural RDM from right pIPS in addition to the skeleton, GBJ, and CorNet-S models. With right pIPS included as a predictor, the skeletal model no longer explained unique variance in left LOC (*β* = 0.07, *p* = .345), only right pIPS (*β* = 0.31, *p* < .001) and CorNet-S (*β* = 0.14, *p* = .053) explained unique variance. By contrast, when linear regression analyses are conducted on right pIPS with the left LOC RDM as a predictor in addition to the skeleton, GBJ, and CorNet-S models, both the skeleton model (*β* = 0.28, *p* < .001) and left LOC RDM (*β* = 0.30, *p* < .001) explain unique variance. Finally, a mediation analysis (with GBJ and CorNet-S as covariates) confirmed that right pIPS fully mediated the relation between the skeletal model and left LOC (*b* = 0.10, 95% CI [.05, .16]). There was no direct relation otherwise (*b* = 0.07, 95% CI [-.074, .21]). By contrast, when left LOC is used as a mediator between the skeletal model and right pIPS, there continues to be a direct relation between the skeletal model and right pIPS (*b* = 0.28, 95% CI [0.14, 0.42]). Here, left LOC acts as only a partial mediator (*b* = 0.05, 95% CI [0.00, 0.10]). Subsequent analyses revealed that other dorsal ROIs (e.g., right aIPS) did not act a mediator between the skeletal model and left LOC. Together these results suggest that object-centered part relations, as approximated by a skeletal model, are computed in right pIPS independently of ventral regions. Moreover, representations of part relation in ventral regions such as left LOC may arise via input from right pIPS.

#### Multivariate connectivity

Thus far, we have documented that an ROI in pIPS, particularly in the right hemisphere, is sensitive to object-centered part relations, able to categorize objects, and account for the representation of part relations in the ventral pathway. Together, these results suggest that this region interacts with ventral regions in support of object recognition. To provide converging evidence for this result, we used multivariate pattern dependence (MVPD) analyses to test whether right pIPS also exhibits functional connectivity with ventral pathway regions during object viewing (see Materials and Methods).

Examination of the group map (Figure 11B) revealed broad connectivity between both right pIPS with bilateral dorsal and ventral regions. To examine the specificity of this interaction between right pIPS and ventral regions, we also examined the multivariate connectivity patterns of left pIPS and bilateral aIPS defined on the basis of part relations. Like right pIPS, these regions also showed broad connectivity with bilateral dorsal and ventral regions (see Figure 11). Direct comparisons between these ROIs (Holm-Bonferroni corrected), revealed that connectivity between right pIPS and bilateral LOC was stronger than both left aIPS (lLOC: *t*(11) = 3.09, *p* = .028, *d* = 0.97; rLOC: *t*(11) = 3.77, *p* = .005, *d* = 1.19) and right aIPS (lLOC: *t*(11) = 2.62, *p* = .072, *d* = 0.83; rLOC: *t*(11) = 3.16, *p* = .019, *d* = 1.00). There were no differences between left and right pIPS (*ps* > .312, *d*s < 0.70), nor among the other ROIs (*ps* > .130, *d*s < 0.71) Together, these findings suggest that right pIPS regions involved in computing object-centered part relations are connected to the ventral pathway.

**Figure 11.**
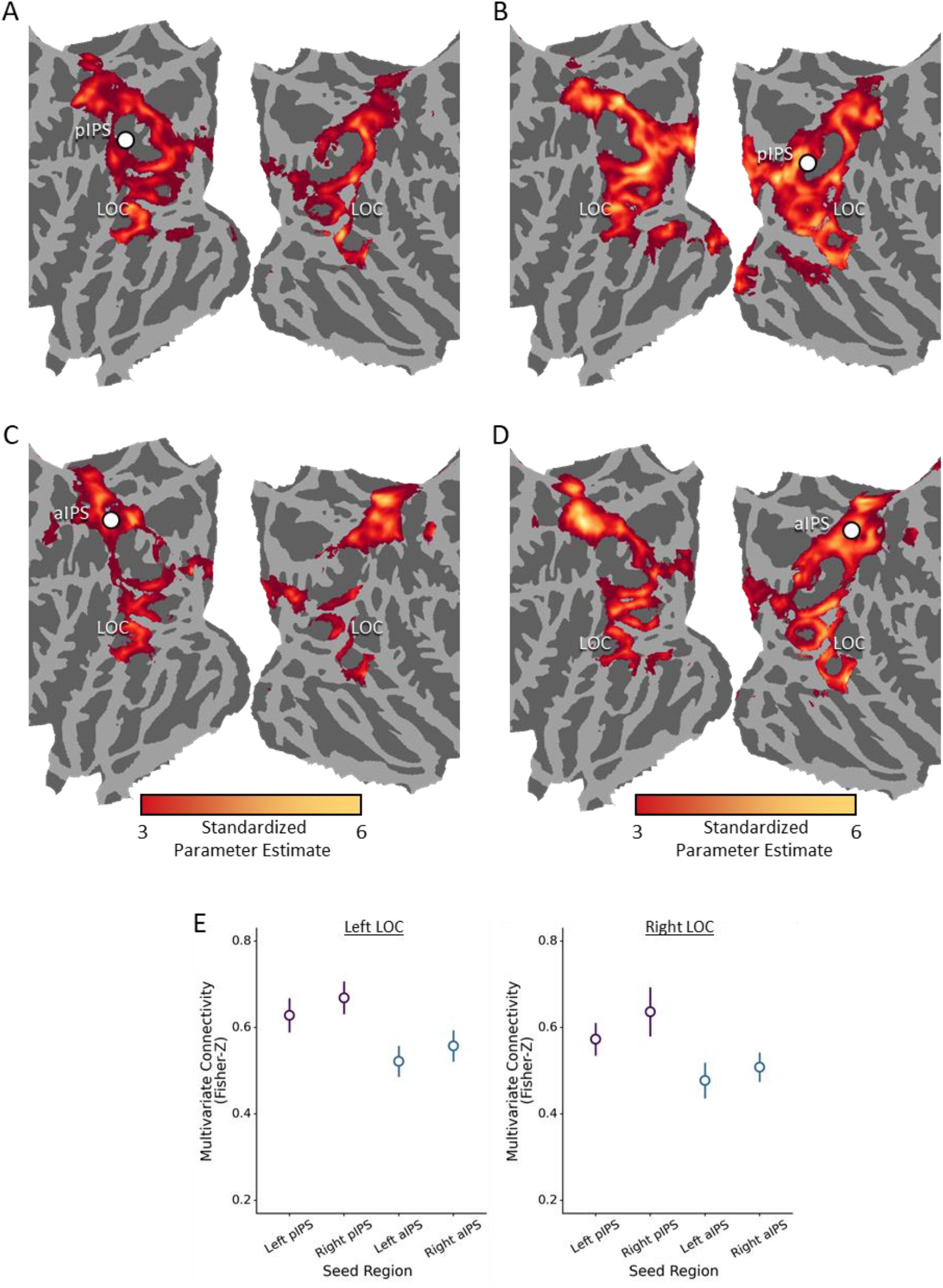
Multivariate functional connectivity results. (A-D) Functional connectivity map for (A) left pIPS, (B) right pIPS, (C) left aIPS, and (D) right aIPS. Seed regions are displayed as a white circle. (C) Plots comparing the connectivity between ROIs in left LOC and right LOC ROIs. Error bars reflect standard error of the mean.

#### Multivariate effective connectivity

If right pIPS transmits information about part relations to LOC for object recognition, then object information should also be processed in right pIPS prior to ventral ROIs. To test this possibility, we conducted multivariate granger causality analyses to test the effective connectivity between IPS regions and LOC (see Materials and Methods).

A Wilcoxon signed-rank comparison to 0 revealed significant effective connectivity between left pIPS with left LOC (*W* = 50, *p* = .010, *r* = 0.82), but not right LOC (*W* = 38, *p* = .161, *r* = 0.38; see Figure 12). Importantly, there was also significant effective connectivity between right pIPS and left LOC (*W* = 61, *p* = .046, *r* = 0.56), as consistent with the mediation analyses presented previously. The effective connectivity between right pIPS and right LOC did not reach significance (*W* = 59, *p* = .065, *r* = 0.51). Finally, there was also significant effective connectivity between left aIPS with left LOC (*W* = 68, *p* = .010, *r* = 0.74), though not right LOC (*W* = 59, *p* = .065, *r* = 0.51), as well as between right aIPS and both left LOC (*W* = 60, *p* = .055, *r* = 0.54) and right LOC (*W* = 71, *p* = .005, *r* = 0.82). Separate repeated-measures ANOVAs were conducted to analyze effective connectivity as function of ROI (pIPS, aIPS) and hemisphere (left, right). These analyses revealed a significant main-effect of hemisphere, such that effective connectivity between right hemisphere IPS ROIs and right LOC were overall higher than left hemisphere IPS ROIs, *F* = 5.37, *p* = .046, η_p_^2^ = 0.37. There were no other significant effects or interactions (*ps* > .451). Thus, as in Experiment 1, these results show that object processing in dorsal cortex precedes and predicts object processing in ventral cortex. Importantly, that pIPS exhibited significant effective connectivity with left LOC is consistent with the hypothesis that pIPS propagates information about part relations to the ventral pathway for object recognition.

**Figure 12.**
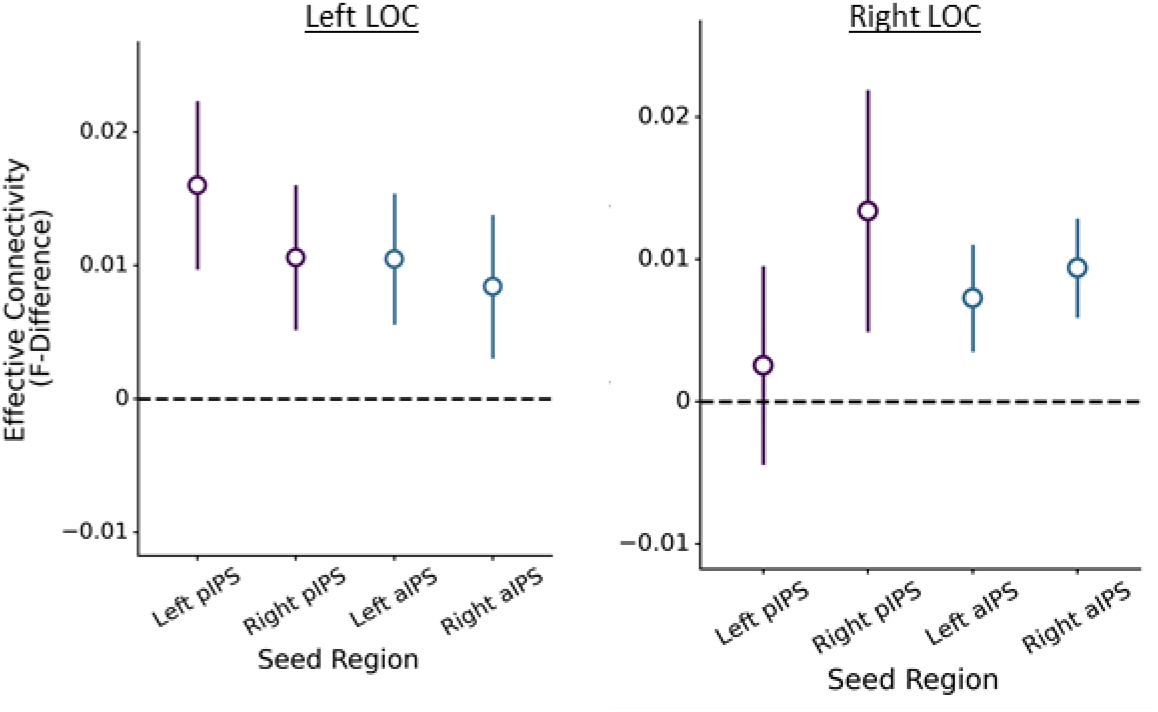
Plots illustrating the multivariate effective connectivity between pIPS and aIPS with left LOC and right LOC ROIs. Error bars reflect standard error of the mean.

#### Analysis on larger sample

All findings from Experiment 2 were replicated successfully with a larger sample (*n* = 14). Object category information was successfully decoded from right pIPS (*p* = .006), as well as left and right LOC (*ps* < .030), but not any of the other ROIs. There was no significant difference in decoding performance between right pIPS with either left or right LOC (*ps* > .264). Representational similarity analyses further showed that objects in right pIPS were best represented by a skeletal model, which approximates the spatial relations among an object’s parts (*p* = .001), rather than the Gabor-jet model or CorNet-S (*ps* > .493). We also found that a skeletal model explained significant variance in left LOC alongside CorNet-S (*ps* < .001). Follow-up analyses revealed that the relation between the skeletal model and left LOC was partially mediated by right pIPS (*p* = .003). Next, multivariate functional connectivity analyses revealed significant functional connectivity between right pIPS with both left and right LOC. In left LOC, this connectivity was significantly greater than left aIPS (*p* < .021), and in right LOC was significantly greater than both left and right aIPS (*ps* < .019). Finally, multivariate effective connectivity analyses revealed significant effective connectivity between right pIPS with both left and right LOC (*ps* < .050).

## General Discussion

Here, we examined the contribution of the dorsal visual pathway to object recognition. Given its sensitivity to spatial information and its contribution to object perception (Freud et al., 2020), we hypothesized that dorsal cortex may compute the relations among an object’s parts and transmit this information to ventral cortex to support object recognition. We found that regions in posterior and anterior IPS, particularly in the right hemisphere, displayed selectivity for part relations independent of allocentric spatial relations and other dorsal object representations, such as 3D shape and tools. Importantly, these regions also exhibited task-dependent functional and effective connectivity with ventral regions, such that connectivity increased when part relations differed.

Next, we found that object category could be decoded successfully in right pIPS, with categorization performance comparable to ventral object regions. Similarity analyses further confirmed that decoding in right pIPS was supported by a representation of part relations, as approximated by a skeletal model, and not by low- or high-level image properties. Crucially, we found that the multivariate response in right pIPS mediated representations of part relations in ventral cortex, with pIPS also exhibiting higher multivariate functional and effective connectivity with ventral cortex. Together, these findings highlight how object-centered part relations, a property crucial for object recognition, are represented neurally, and validate the strong link between dorsal and ventral visual cortex in accomplishing object recognition.

### Neural representations of object-centered part relations

Many studies have examined how allocentric spatial information is represented neurally, but few have explored the representations of object-centered part relations. Lescroart and Biederman (2012) decoded the spatial arrangements of object parts in both ventral and dorsal cortices, but did not test whether these were independent of other dorsal representations nor whether other visual properties influenced decoding. Ayzenberg et al. (2021) identified ventral regions that coded for part relations (as approximated by a skeletal model) independent of other visual properties, with strongest coding in left LOC – a finding consistent with the RSA results of the current study. However, they did not investigate whether such representations also exist in dorsal cortex and could account for their effects. Finally, Behrmann et al. (2006) reported that patients with LOC damage and object recognition deficits were impaired in perceiving part relations, but not the features of object parts, suggesting a ventral locus for object-centered relations.

Consistent with these studies, we, too, found that part relations are represented in ventral cortex. However, our data suggest that this information arises via input from dorsal cortex. We documented functional connectivity between IPS and LOC, and showed that right pIPS mediates the representation of part relations in ventral regions, and not the other way around. Indeed, across both experiments, effective connectivity analyses revealed that part relations may be first processed in IPS and then transmitted to ventral object regions. This finding is compatible with research showing that visual object information reaches posterior parietal cortex 100 to 200 ms earlier than ventral regions (Regev et al., 2018), as well as with studies showing that topological object properties may only become represented in the ventral pathway through top-down connections (Bar et al., 2006; Wang et al., 2020). Crucially, studies also show that temporary inactivation of posterior parietal regions impairs ventral object processing (Van Dromme et al., 2016; Zachariou et al., 2017). Altogether, our results in combination with these studies suggest a causal role for dorsal cortex in ventral object processing in which dorsal cortex transmits object information to the ventral pathway to support object recognition.

An interesting facet of our work is that our results differed by hemisphere. Specifically, we found that coding of object-centered part relations was strongest in the right hemisphere across almost all analyses. This finding mirrors the classic global precedence effect of the right hemisphere (Brighina et al., 2003; Van Kleeck, 1989; Wasserstein et al., 1987), wherein global shape properties are most often represented by the right hemisphere and local shape properties by the left. Although the reasons for this effect remain controversial (Kimchi, 1992; Seghier & Vuilleumier, 2006), one explanation suggests that the right hemisphere may be more sensitive to low spatial frequencies (Iidaka et al., 2004; Peyrin et al., 2004). Consistent with this possibility patients with damage to posterior parietal cortex show a deficit in perceiving low spatial frequency information, and, as a result, global form (Kinsbourne & Warrington, 1962; Thomas et al., 2012; Warrington & Taylor, 1973). Other studies suggest that the right hemisphere global precedence may be related to lateralization of object-based attention to the right hemisphere (Shomstein & Behrmann, 2006), such that manipulating the focus of attention can enhance or disrupt the global precedence effect in the right hemisphere (Kimchi & Merhav, 1991; Van Vleet et al., 2011).

Our results also uncovered a posterior-to-anterior gradient, especially evident in Experiment 2. Although selectivity for part relations was found in both pIPS and aIPS, only right pIPS was able to decode object category. Moreover, right pIPS exhibited the highest multivariate functional connectivity with LOC, and its representation of object similarity was most consistent with a model of part relations (i.e., medial axis skeleton). This gradient may reflect a common organizing principle of the dorsal pathway. Regions of posterior parietal cortex exhibit greater sensitivity for object properties in the service of recognition (Freud et al., 2017; Gillebert et al., 2015; Van Dromme et al., 2016), and greater connectivity to ventral object regions (Janssen et al., 2018; Takemura et al., 2016; Webster et al., 1994). By contrast, anterior parietal cortex shows greater sensitivity to object properties that afford action, such as elongated axes (Chao & Martin, 2000; Chen et al., 2017; Chen et al., 2016; Culham et al., 2003). Whereas right pIPS may be more involved in computing part relations for the purpose of recognition, right aIPS may be more involved in computing part relations to help coordinate grasping behaviors. Relatedly, we found greater overlap between right aIPS and regions involved in representing allocentric relations and tools – which are both critical for coordinating action. However, it is important to note that right aIPS did show significant functional and effective connectivity with ventral regions. Given the research described above, it is possible that that right aIPS may contribute to categorization for objects that afford action, such as tools. Unfortunately, none of the objects used in Experiment 2 consisted of tools, and only two of the object categories (out of five) could be considered manipulable. Thus, future research should explore the degree to which dorsal cortex may differentially contribute to object categorization for manipulable and non-manipulable objects.

### Object-centered relations and other dorsal representations

We found that IPS regions responded more to object-centered part relations than allocentric relations, 3D shape, and tools, suggesting selectivity in these regions. However, our conjunction analyses also revealed that object-centered relations may be represented along a continuum in parietal cortex, with varying degrees of overlap with other dorsal properties, particularly, with allocentric spatial relations. The overlap between object-centered and allocentric relations in parietal cortex may reflect a broader organizing principle for spatial coding in dorsal cortex in which reference frames are organized topographically. Recent evidence suggests that the dorsal pathway represents visual information at different spatial scales ranging from single objects to large, multi-object perspectives (Josephs & Konkle, 2020). This possibility is also consistent with a rich literature on hemi-spatial neglect, in which right parietal damage impairs object perception on the left side of space (Caramazza & Hillis, 1990; Corbetta & Shulman, 2011; Heilman & Valenstein, 1979). Depending on the scope of the damage, multiple reference frames are often affected simultaneously, further suggesting that the representations overlap or abut (Halligan et al., 2003; Medina et al., 2009). However, our data is also consistent with studies showing distinct representations of object-centered reference frames (Vannuscorps et al., 2021a; Vannuscorps et al., 2021b). These representations are crucial for object perception and are most likely mediated by the dorsal pathway (Freud & Ahsan, 2022; Taylor & Xu, 2022). Altogether, we suggest that such representations are situated within a broader topographic map for spatial coding.

We found relatively little overlap between regions involved in representing part relations and those involved in representing tools – with overlap occurring exclusively in aIPS. This finding is consistent with the hypothesis formulated earlier, that coding of part relations in aIPS may be in support of coordinating grasping behaviors. It is important to note that here we used a particularly stringent definition of tool ROIs, wherein tools were contrasted with other manipulable objects (Chen et al., 2018), and this decision may have minimized the degree to which we observed activity related to object action affordances (since all stimuli afforded action). Moreover, by using objects with elongated axes in the part-relations localizer (an important indicator of action affordance; Chen et al., 2017), we may have further suppressed the degree to which regions representing part relations overlapped with those representing tools. Future work may use a more direct object affordance localizer (Freud et al., 2018; Snow et al., 2011) and a more variable stimulus set to localize part relations.

Finally, extensive pilot work (Ayzenberg et al., unpublished data) suggested that depth regions in parietal cortex could be reliably localized with the 3D and 2D shape stimuli used here. However, we were unable to do so in current study – precluding conjunction analyses. Two runs of the depth localizer may have been insufficient to identify regions involved in processing 3D shape, and/or depth from shading (as used here) may be less consistently represented than depth from texture or disparity (Georgieva et al., 2008). Given that the computation of depth structure in the dorsal pathway is critical for object recognition (Farivar, 2009; Freud et al., 2020; Van Dromme et al., 2016; Welchman, 2016), future work is required to explore the link between regions subserving part relations and 3D shape.

### The role of object-centered part relations in object recognition

Representations of object-centered part relations are thought to be critical for object recognition because they describe an object’s global shape structure – a key organizing feature of most basic-level categories (Barenholtz & Tarr, 2006; Hummel, 2000; Mervis & Rosch, 1981). Such a representation may even support rapid object learning in infancy when experience with objects is minimal (Ayzenberg & Lourenco, 2021; Kraebel & Gerhardstein, 2006; Rakison & Butterworth, 1998). Yet, ANNs, the current best models of human object recognition, are largely insensitive to the relations among object parts and require extensive object experience to categorize novel objects (Baker et al., 2018; Baker et al., 2020). One potential reason for this deficit is that most current ANNs exclusively model ventral cortex processes (Blauch et al., 2021; Schrimpf et al., 2020; Yamins et al., 2014). Indeed, the few ANNs that model dorsal cortex focus on action or motion related processes (Güçlü & van Gerven, 2017; Mineault et al., 2021). Here, we propose that the dorsal pathway may play a key role in object recognition by computing object-centered part relations and propagating these signals to ventral object regions. Right pIPS, in particular, may be important for object recognition, in that its multivariate response was sufficient to decode object category and it was well explained by an object recognition model that computes part relations (i.e., a skeletal model). Importantly, we consistently found connectivity between right pIPS regions and regions in ventral cortex, with evidence that right pIPS may even mediate the representation of part relations in LOC. Thus, by incorporating the dorsal pathway with the ventral pathway, we may gain a better understanding of the broader network that supports object recognition and the relative contributions of each pathway.

## Acknowledgments and funding

This work was supported by a National Science Foundation (NSF) grant (BCS2123069) awarded to M.B.

## Data availability

Data, stimuli, and tasks are available at: https://doi.org/10.1184/R1/19543819.v1

## Code availability

Analysis and modelling scripts are available at: https://github.com/vayzenb/dorsal-part-relations

